# Protein stability is determined by single-site bias rather than pairwise covariance

**DOI:** 10.1101/2025.01.09.632118

**Authors:** Matt Sternke, Katherine W. Tripp, Doug Barrick

## Abstract

The biases revealed in protein sequence alignments have been shown to provide information related to protein structure, stability, and function. For example, sequence biases at individual positions can be used to design consensus proteins that are often more stable than naturally occurring counterparts. Likewise, correlations between pairs of residue can be used to predict protein structures. Recent work using Potts models show that together, single-site biases and pair correlations lead to improved predictions of protein fitness, activity, and stability. Here we use a Potts model to design groups of protein sequences with different amounts of single-site biases and pair correlations, and determine the thermodynamic stabilities of a representative set of sequences from each group. Surprisingly, sequences excluding pair correlations maximize stability, whereas sequences that maximize pair correlations are less stable, suggesting that pair correlations contribute to another aspect of protein fitness. Consistent with this interpretation, we find that for adenylate kinase, enzyme activity is greatly increased by maximizing pair correlations. The finding that elimination of covariant residue pairs increases protein stability suggests a route to enhance stability of designed proteins; indeed, this strategy produces hyperstable homeodomain and adenylate kinase proteins that retain significant activity.

**Significance statement:** Recent methods for protein structure analysis and design have used sequence covariance to help predict protein structure, stability, and function. Here, by designing homeodomain and adenylate kinase sequences with different amounts of single-site bias and pairwise covariance, we find that stability is solely determined by single-site bias but not pairwise covariance. However, pairwise covariance makes an important contribution to catalysis in adenylate kinase. Our findings suggest a new way to generate highly stable proteins: by separating single-site biases from pairwise covariance, the single-site coefficients can be used to design proteins with stabilities even higher than those obtained by consensus design.

## Introduction

Most proteins fold into stable, well-defined native structures and perform specific biological functions. Over evolutionary time, mutations introduce random amino acid substitutions which are subject to physiochemical constraints that retain native-state structure and biological function. Because protein structure and function are determined by the many cooperative interactions among their amino acids,^1^ these residue-residue interactions play a role in shaping and constraining protein sequence evolution, and lead to statistical correlations among pairs of residues within protein sequences.

Residue covariance has been shown to be important for specifying protein structure and function. Using an approach called statistical coupling analysis (SCA), Ranganathan and coworkers demonstrated that maintaining residue covariances found in a multiple sequence alignment (MSA) is necessary for proper folding of WW domains.^2^ Subsequent studies showed that SCA-based protein designs retain expected biological activity^3,4^ More recently, a complementary approach using residue co-evolution called direct coupling analysis (DCA) has gained popularity for analyzing protein structure and function. DCA uses a Potts formalism from statistical mechanics to separate position-specific single-site biases from pairwise coupling biases.^5,6^ Starting with an MSA, the Potts model infers single-site and pairwise coupling energy coefficients. Including both single-site biases and pair correlations from the Potts formalism has been shown to improve predictions of protein structure^5^, predictions of effects of mutations on protein stability and function^7,8^, and have been used to design non-natural sequences that retain biological function.^9,10^

Although correlations between pairs of residues have been shown to contribute to protein structure and function, design strategies that do not explicitly include these correlations have been quite successful. These strategies include ancestral reconstruction and consensus design, which infer residues independently at each site without considering interactions between residues.^11^ ^12^ Despite this potentially naïve assumption of site-independence, both strategies have widely demonstrated success in producing folded, stable, and biologically-active proteins with sequences.^13–17^ In many cases, ancestral and consensus proteins show greater stabilities than natural proteins, highlighting the effectiveness of site-independent models in capturing information specifying protein stability.^18,19^

To better understand the contributions of sequence covariance to protein stability and function, and to explore why single-site models achieve high levels of success in protein design while ignoring covariance, we used a Potts model to generate and analyze a large set of sequences for two well-studied families: the DNA-binding homeodomain family and the adenylate kinase enzyme family. For each family we designed sequences that differ in the relative amounts of pairwise coupling and single-site bias, allowing us compare the effects of these two biases to protein stability and activity.

## Results

### Potts model inference and sequence design for homeodomains

To determine single-site and coupling energy coefficients (*h_i_(a)* and *j_i,k_(a,b)*, where *i* and *k* indicate positions and *a* and *b* indicate amino acid types) we fit a Potts model to a large alignment of HD sequences (Fig 1; see Methods). This fitting procedure produces *h_i_(a)* coefficients for all residues at all positions and *j_i,k_(a,b)* coefficients for all pairs of residues at all pairs of positions Using these Potts coefficients, we generated energy functions that include different amounts of single-site and pairwise coupling energies. These include an energy function that includes all of the pairwise coupling energies along with the single-site energies (*ɛ*(𝑠𝑒𝑞)_𝐻𝐽_, equation 6) and an energy function that omits pairwise terms, using only single-site energies (*ɛ*(𝑠𝑒𝑞)_𝐻_, equation 9).

**Figure 1.**
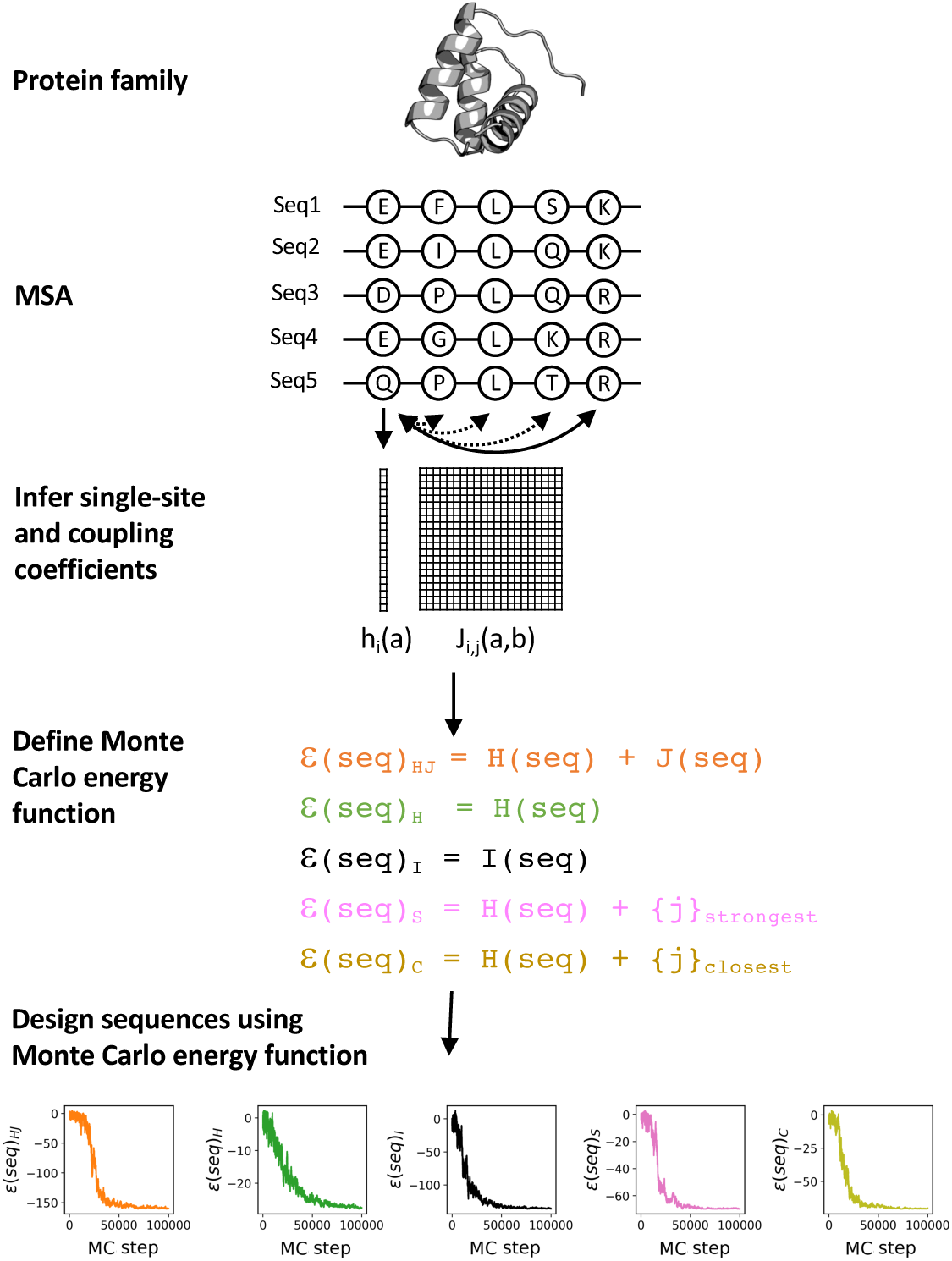
Potts-based design of protein sequences. Single-site and pairwise coupling energies are inferred from an MSA using a Potts model (here the homeodomain family, PDB: 1ENH). Energy functions are created using different amounts of single-site versus pairwise energies in the sequence design, and are used to generate sequences with a Monte Carlo search. These sequences, which contain different relative amounts of intrinsic and pairwise coupling energy, are characterized for stability and activity.

In addition, we fit the same HD MSA with a model that infers only single-site energies, ignoring all pairwise couplings (Figure 1, equations 12 and 13). The single-site energies (*I_i_(a)*) inferred from this model, which are closely related to the marginal residue frequencies in the MSA were used to construct an energy model (*ɛ*(𝑠𝑒𝑞)_𝐼_) for generating sequences based solely in single-site conservation. Sequences designed using *ɛ*(𝑠𝑒𝑞)_𝐼_ are closely related to consensus sequences (indeed, the designed HD sequence with the lowest *I(seq)* score is identical to the consensus sequence for the MSA used here).

For each of these energy functions, we used a Monte Carlo search procedure to generate 1,000 independent low-energy homeodomain sequences. The sequences generated from these energy models show features that are consistent with the energy functions used to create them. Sequences designed with *ɛ*(𝑠𝑒𝑞)_𝐻_, referred to as H-optimized sequences, have high *H(seq)* values but low *J(seq)* values (green, Figure 2), whereas sequences designed with *ɛ*(𝑠𝑒𝑞)_𝐻𝐽_, referred to as HJ-optimized sequences, have high *J(seq)* values but low *H(seq)* values. Although *h_i_(a)* and *j_i,k_(a,b)* are equally weighted in *ɛ*(𝑠𝑒𝑞)_𝐻𝐽_, there are many more *j_i,k_(a,b)* terms in *J(seq)* than there are *h_i_(a)* terms in *H(seq)* (21^2^*×L(L-1)/2* versus *21L* for a sequence of length *L* residues). Thus, HJ-optimization is dominated by the *j_i,k_(a,b)* coupling energies (for the HJ-optimized HD sequences the total *J(seq)* values are about seven times larger than the total *H(seq)* values; Figure 2). As a result, HJ-optimized sequences are quite similar to sequences optimized with *j_i,k_(a,b)* terms alone.

**Figure 2.**
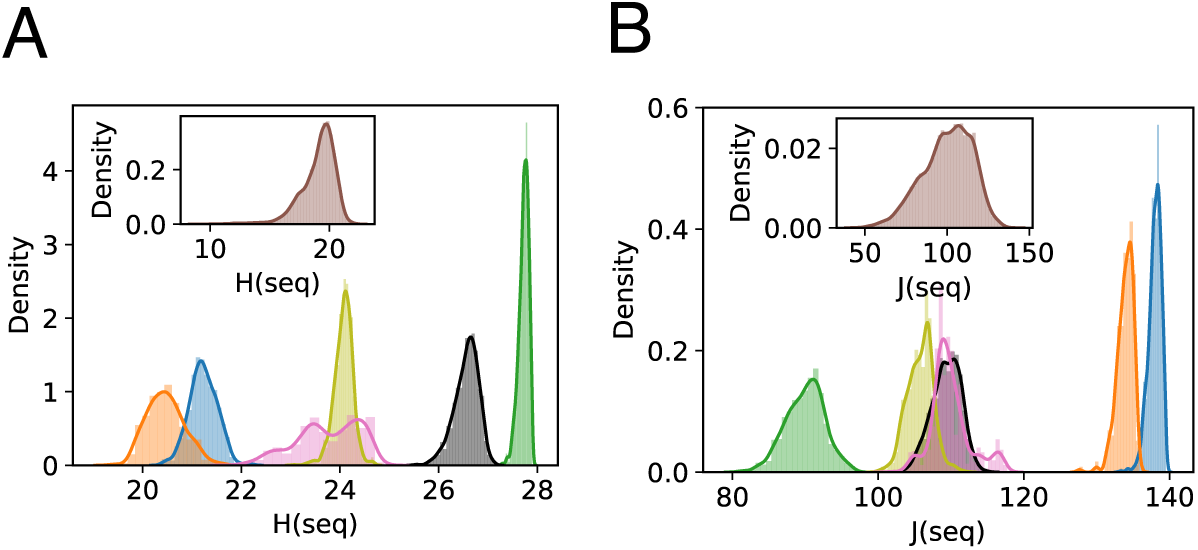
Sequence design of HDs. Distributions of total single-site energies (A) and total pairwise coupling energies (B) for the 1,000 sequences optimized using different energy functions. Green: *ɛ*(𝑠𝑒𝑞)_𝐻_; orange and blue: *ɛ*(𝑠𝑒𝑞)_𝐻𝐽_; grey, *ɛ*(𝑠𝑒𝑞)_𝐼_; pink: *ɛ*(𝑠𝑒𝑞)_𝐻𝐽𝑆_; yellow: *ɛ*(𝑠𝑒𝑞)_𝐻𝐽𝐶_. Insets show distributions for the 19,221 extant HDs in the MSA.

Interestingly, HJ-optimized sequences have a bimodal distribution of *H(seq)* and *J(seq)* values (Figure 2), suggesting that designing sequences with high pairwise coupling energies captures higher-order sequence correlations across three or more sites. Consistent with this, pairwise identities among the 1000 HJ-optimized sequences have a bimodal distribution (Figure S1). This distribution of identities results from two distinct clusters of sequences of roughly equal size, which we designate as clusters A and B. Within clusters A and B, sequences have high identities (89 and 88 percent respectively, Figure S1), whereas between clusters sequences have lower identity (55 percent). In contrast, sequences designed using the *ɛ*(𝑠𝑒𝑞)_𝐻_ energy function have unimodal *H(seq)*, *J(seq)*, and sequence identity distributions (Figures 2, S1).

Comparison of sequences generated using the independent model (with the *ɛ*(𝑠𝑒𝑞)_𝐼_ energy function, referred to as I-optimized sequences) to those generated from the Potts model reveal some subtle differences. Although the *H(seq)* values of the I-optimized sequences are high, as might be expected for a design strategy based on single-site frequencies, values are not as high as those of the H-optimized sequences (grey and green, Figure 2A). Likewise, the *J(seq)* values of the I-optimized sequences are low, but they are not as low as those of the H-optimized sequences (Figure 2B). Rather, their *J(seq)* values are midway between those of the H-optimized and HJ-optimized sequences. These differences are reflected in sequence identities (Figure S1): the I-optimized sequences are closer to both the H-optimized sequences (78 percent average pairwise identity) and HJ-optimized sequences (63 and 58 percent identity to clusters A and B) than the H-optimized and HJ-optimized sequences are to one another (53 percent identity to both clusters A and B). The finding that I-optimized sequences have *J(seq)* values midway between the H- and HJ-optimized sequences demonstrates that I-optimization inadvertently introduces pairs of residues that are positively covariant, which might be expected for covariant pairs that are strongly conserved. This observation also indicates that sequences designed using *ɛ*(𝑠𝑒𝑞)_𝐻_ are purged of pairs of correlated residues. Thus, along with the HJ-optimized sequences, the H-optimized sequences provide a stringent test set to evaluate the relative effects of single-site biases and pair correlations on protein stability and function, maximizing *H(seq)* values and minimizing *J(seq)* values beyond what is obtained with commonly used single-site strategies such as consensus design.

### Biophysical characterization of designed and extant HD proteins

To examine how single-site and pairwise coupling energies influence stability and biological activity, we selected proteins from each Monte Carlo optimization to express, purify, and characterize experimentally (Table S1). Because each Monte Carlo sequence optimization generates some sequence variation (Figure S1), we characterized multiple proteins for each optimization. For each optimization, we picked the sequence with lowest *ɛ*(𝑠𝑒𝑞) value (which we refer to as Seq1), the sequence with the mean *ɛ*(𝑠𝑒𝑞) value (Seq3), the sequence with an *ɛ*(𝑠𝑒𝑞) value one standard deviation below the mean (Seq2), and the sequence with an *ɛ*(𝑠𝑒𝑞) value one standard deviation above the mean (Seq4). Thus for each optimization, *ɛ*(𝑠𝑒𝑞) values increase monotonically from Seq1 to Seq4. For the HJ-optimized sequences, we characterized four such proteins from each of the A and B clusters. The sequences selected from each Monte Carlo sequence optimization have in-group identities ranging from 65% to 96% (Fig S3). To attempt to provide an unbiased representation of the biophysical properties of extant HD proteins, we also expressed and characterized five proteins from the MSA as well as the Engrailed HD from *D. melanogaster* used in our previous study (Table S1).^15^ These extant sequences share 26% to 53% identity to one another (Fig S3). All of these extant homeodomains expressed in *E. coli*; however, one of these proteins displayed limited solubility and was not characterized biophysically. Sequences are referred to as X_Seq#, where X indicates the energy function used for sequence generation (H for H-optimization, HJA and HJB for HJ-optimization cluster A and B, I for independent sequence optimization, and E for extant protein).

Most of the Monte Carlo designed HD proteins expressed and were soluble. However, three out of four sequences from HJ cluster 2 had limited solubilities and could not be characterized. For all remaining proteins, far-UV CD spectra show minima at 208 and 222 nm that are similar in magnitude to those of extant HDs, indicating that all designed proteins adopt stable α-helical structures (Fig S3). To examine how the single-site and pairwise coupling energies contribute to stability, we collected GdnHCl-induced unfolding transitions for the designed and extant HD proteins (Fig 3A). All proteins show sigmoidal unfolding transitions with similar slopes indicating that the designed HDs retain similar folding cooperativity as the extant HDs. Extant HDs have folding free energies ranging from -2.38 to -5.45 kcal mol^-1^ with a mean of -3.91 kcal mol^-1^ (Fig 3B, Table S2). HD sequences from the *ɛ*(𝑠𝑒𝑞)_𝐻𝐽_ optimization are more stable than the extant HDs, with sequences from cluster A having folding free energies ranging from -6.84 to -9.36 kcal mol^-1^ with a mean of -8.20 kcal mol^-1^; the single sequence we were able to purify from cluster B (HJB_Seq1) has a folding free energy of -6.45 kcal mol^-1^. (Fig 3B, Table S2). Thus, including both the single-site and pairwise coupling information from the MSA increases protein stabilities compared to the extant sequence collection (by an average ΔΔG of -4.29 and -2.54 kcal mol^-1^ for cluster A and HJB_Seq3, Figure 3B). This result is consistent with previous studies that show correlation between protein stability and Potts energies determined by equally weighting single-site and pairwise coupling information.

**Figure 3.**
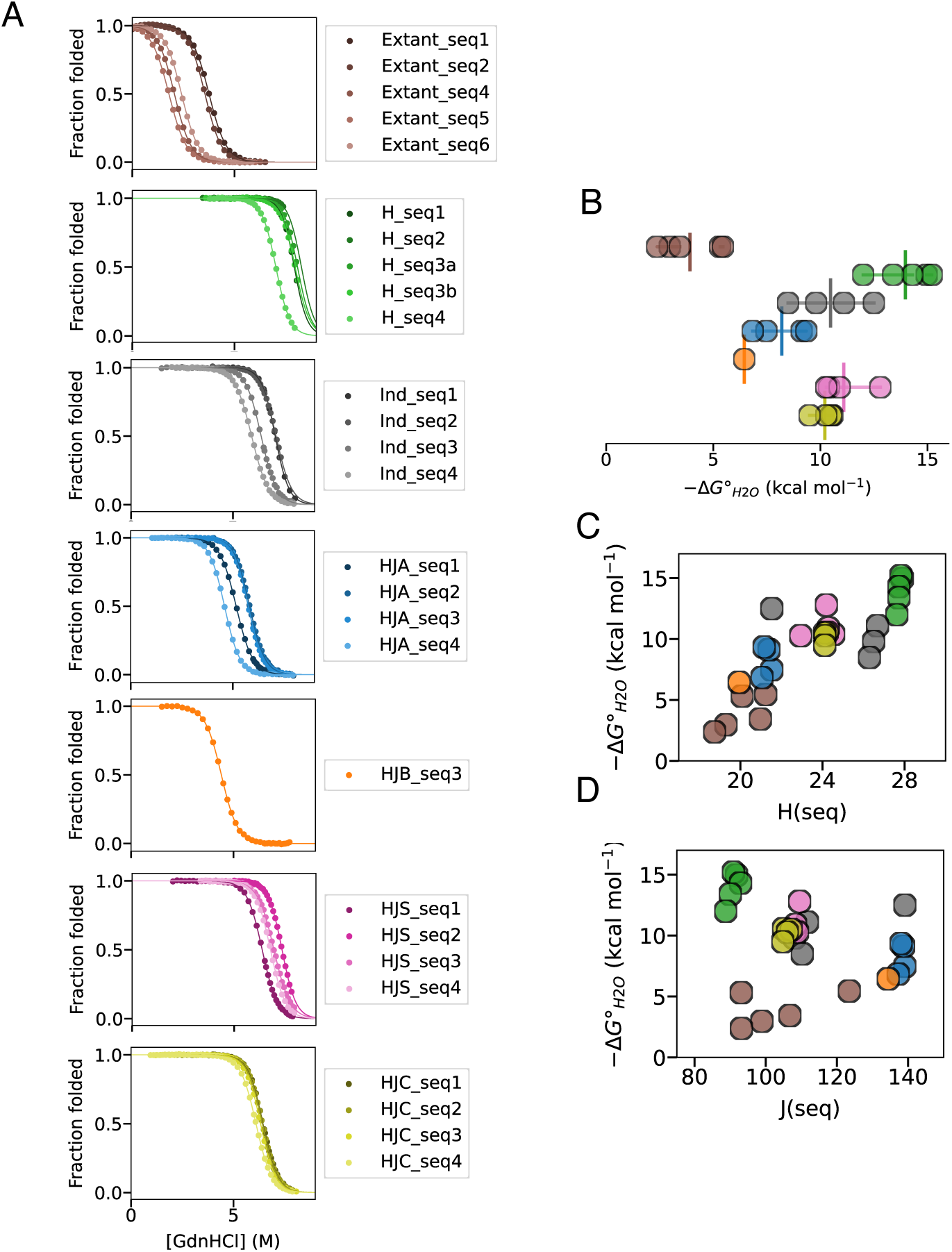
Stabilities of Potts-designed HDs. (A) Representative GdnHCl-induced unfolding transitions of HDs at 25 °C. Solid lines represent the fit of a two-state unfolding model; for sequences with incomplete high-temperature baselines, fits included additional GdnHCl transitions at higher temperatures (Figure S4). (B) Folding free energies of HDs determined from the two-state unfolding analyses as in panel B. Vertical bar indicates the mean folding free energy of each distribution. (C and D) Correlation of folding free energies and sequence single-site energies (C) and sequence coupling energies (D). Sequences were generated with a sequence reweighting X_ID_=0.8 and regularization parameters λ_h_=λ_j_=0.01.

To determine how the relative contributions of the single-site and pairwise coupling energies to protein stability, we measured GdnHCl-induced unfolding transitions of HD sequences generated by optimizing *ɛ*(𝑠𝑒𝑞)_𝐻_. To our surprise, these H-optimized sequences are significantly more stable than those generated by HJ-optimization. In fact, the sampled H-optimized sequences were so stable that the unfolded baseline was not resolved at 25 °C (Figure 3A). For these proteins, stabilities at 25 °C had to be determined using a global analysis including unfolding transitions at elevated temperatures (see Methods, Fig S4). Folding free energies of H-optimized sequences at 25 °C ranged from -12.0 to -15.2 kcal mol^-1^, with a mean of -14.0 kcal mol^-1^. This observation does not support the idea that increases in stability of Potts designed sequences result from optimizing pairwise sequence energies (the HJ-optimized sequences have much larger *J(seq)* values than the H-optimized sequences, Figure 2, but are significantly less stable). Rather, it suggests that the stability increase results from optimizing single-site energies (the H-optimized sequences have much larger *H(seq)* values than the HJ-optimized sequences, Figure 2); indeed, a positive correlation is seen between stability and *H(seq)* values for all of the HD sequences examined here (Potts, independent, and extant; Figure 3C). Alternatively, it is possible that positively covariant residue pairs are destabilizing, although analysis with a larger set of sequences (see below) does not support such a correlation.

The magnitude of the increase in protein stability for the H-optimized sequences is large, especially considering the small size of the Homeodomain sequences (57 residues). The average folding free energy of the H-optimized sequences increases by -10.1 kcal mol^-1^ compared to the average for extant HD sequences (-13.4 versus -3.9 kcal mol^-1^, Figure 3B). This is a considerably larger stability increase than we saw previously for consensus homeodomains designed from a smaller sequence alignment. To compare to stabilities of a consensus-like sequences designed from the same alignment and Monte Carlo design methods used here for Potts analysis, we measured stabilities of sequences designed using the *ɛ*(𝑠𝑒𝑞)_𝐼_ energy function. The *I_i_(a)* coefficients in this energy function are generated from local biases without consideration of pair correlations, as is also the case for consensus design.

These *ɛ*(𝑠𝑒𝑞)_𝐼_-optimized HDs have folding free energies ranging from -8.49 to -12.5 kcal mol^-1^ with a mean of -10.5 kcal mol^-1^ (Fig 3C, Table S2). On average, the H-optimized sequences are more stable than the *ɛ*(𝑠𝑒𝑞)_𝐼_-optimized sequences by a ΔΔG° value of -3.5 kcal mol^-1^. A similar stability increment

(ΔΔG° = -2.7 kcal mol^-1^) is seen between the most stable H-optimized sequence (H_Seq2) and the most stable I-optimized sequence (the consensus, Ind_Seq1). Thus, at least for HD sequences, it appears that designing sequences using H-optimized coefficients can substantially increase stability beyond that obtained by consensus design.

In contrast, the I-optimized sequences are more stable than the HJ-optimized sequences by an average ΔΔG° value of -2.2 kcal mol^-1^ (cluster A) and -4.0 kcal mol^-1^ (cluster B HJB_Seq3). This increase in stability for the consensus-like sequences generated from the independent model may be due to the larger *H(seq)* energies for the I-optimized versus HJ-optimized sequences (Figure 2A).

### Stabilities of HD sequences with restricted pairwise coupling energies

Our finding that HD sequences designed using pairwise coupling information are less stable than sequences designed using single-site bias (either through H-optimization or consensus-like I-optimization) is unexpected, given previous work demonstrating the importance of residue couplings in promoting folding and predicting structural contacts.^2,5^ One possible explanation for this finding is that optimization using pair coupling terms is dominated by a large number of noisy *j_i,k_(a,b)* terms from uncoupled sites. To mitigate the effects of potential noise that may be introduced by small coupling terms, we designed sequences using energy functions in which small coupling terms are omitted. In one design, we optimized sequences using an energy function, *ɛ*(𝑠𝑒𝑞)_𝑆_, in which coupling terms are restricted to the ten percent of residue pairs that have the strongest couplings (see Methods). In another design, we optimized sequences using an energy function, *ɛ*(𝑠𝑒𝑞)_𝐶_, in which coupling terms are restricted to the ten percent of residue pairs that are closest in the *D. melanogaster* Engrailed HD structure (PDB: 2JWT). These two subsets of pair positions are similar (but not identical, Figure S9A, right), consistent with the idea that strongly coupled residue pairs are close in three dimensional structure^20^.

Using the *ɛ*(𝑠𝑒𝑞)_𝑆_ and *ɛ*(𝑠𝑒𝑞)_𝐶_ energy functions, we designed HJS- and HJC-optimized HD sequences using the Monte Carlo approach described above (Table S1) . These two sets have *H(seq)* and *J(seq)* values midway between HJ and H-optimized sequences (Figure 2). HJS- and HJC-optimized sets of proteins are similar in stability (Figure 3, mean ΔG° values of -11.1 and -10.2 kcal mol^-1^ respectively, Figure 3). The HJS- and HJC-optimized sequences are more stable than the HJ-optimized sequences (mean ΔΔG° values are -2.9 and -2.0 kcal mol^-1^), but are less stable than H-optimized sequences (mean ΔΔG° values are +2.9 and +3.8 kcal mol^-1^). These results suggest that the lower stabilities of the HJ-optimized sequences compared to the H-optimized sequences are not the result of noise introduced from weak coupling coefficients, but from the stabilizing effects of single-site *h_i_(a)* energy terms and the potential destabilizing effects of pairwise *j_i,k_(a,b)* energy terms.

### Sensitivity of HD stabilities to global Potts fitting parameters

Fitted values of Potts *h_i_(a)* and *j_i,k_(a,b)* terms can be affected by three global parameters: two regularization parameters (λ_h_ and λ_j_) and a sequence reweighting parameter (X_ID_). The regularization parameters help prevent overfitting of the *h_i_(a)* and the many *j_i,k_(a,b)* terms (∼700,000 for homeodomain), most of which correspond to unobserved or infrequently observed residue pairs and are thus poorly defined by the MSA. The sequence weighting parameter down-weights highly similar sequences in the MSA to avoid taxonomic sequencing biases. For the HD sequences described above, we used the parameters λ_h_=λ_j_ =0.01 and X_ID_=0.8, which have been used in previous Potts analyses of protein families^21^. To test whether the findings above are dependent on the values we chose for regularization and sequence reweighting, we generated a set of HD sequences using the parameters λ_h_=λ_j_=0.1 and X_ID_=0.2. Increasing the regularization parameters by a factor of 10 should decrease potential noise in the fitted *h_i_(a)* and *j_i,k_(a,b)* coefficients but may bias these coefficients to small values . Decreasing the sequence reweighting parameter should result in a more uniform weighting, since nearly all sequences are identical to one another at the 20 percent threshold (Figure S1).

Using this new set of global parameters, Potts coefficients were inferred from the same HD MSA used for the analysis above. These coefficients are similar to those determined in the previous section. In both cases, the two sets of coefficients are linearly related, with Pearson correlation coefficients of 0.94 and 0.65 respectively (Figure S5). As above, these Potts coefficients were combined in different proportions into energy functions for Monte Carlo sequence generation (Table S2). These included an energy function using only *H(seq)* values (*ɛ*(𝑠𝑒𝑞)_𝐻_, equation 9), a function equally weighting *H(seq)* and *J(seq)* values (*ɛ*(𝑠𝑒𝑞)_𝐻𝐽_, which again generated two clusters of sequences), and a function that down-weights *J(seq)* by a factor of 20 so that it is equal in magnitude to *H(seq)* (*ɛ*(𝑠𝑒𝑞)_𝐻+0.05𝐽_).

Overall, the HD sequences designed with the global parameters X_ID_=0.2, λ_h_=λ_j_=0.1 show the same stability patterns as those described above (Figure S6, Table S4). The HJ sequences optimized with the global parameters X_ID_=0.2, λ_h_=λ_j_=0.1 are more stable than extant sequences (by an average ΔΔG of - 2.4 and -3.6 kcal mol^-1^ for cluster A and B), although they are considerably less stable than H-optimized sequences (by an average ΔΔG of +5.3 and +4.1 kcal mol^-1^). On average, the H sequences optimized with X_ID_=0.2, λ_h_=λ_j_=0.1 are more stable than the consensus (I-optimized) sequence, (also generated with X_ID_=0.2) by an average ΔΔG of -1.3 kcal mol^-1^, and the most stable H-optimized sequence is more stable than consensus by -4.2 kcal mol^-1^. These trends are consistent with stability increases in Potts constructs result from optimizing *H(seq)* rather than *J(seq)*. Consistent with this observation, sequences optimized from the X_ID_=0.2, λ_h_=λ_j_=0.1 Potts coefficients where *J(seq)* is damped 20-fold (so that *H(seq)* and *J(seq)* values are weighted equally in the Monte Carlo search) have folding free energies midway between the H- and HJ-optimized sequences (Figure S6, Table S4).

### Correlation between HD stability, single-site, and pairwise coupling energies

Together, the HD sequences characterized here from the various Potts design strategies (Tables S1, S2, S5; 62 sequences total including extant sequences) provide a large data set to evaluate how stability is related to the *H(seq)* and *J(seq)* scores. Using the negative of the folding free energy as a measure of stability, we find a moderate positive correlation between −Δ𝐺°_𝐻20_values and *H(seq)* values (calculated with the Potts coefficients obtained using global parameters X_ID_=0.8. λ_h_=λ_j_=0.01), with a Pearson correlation coefficient of 𝜌_𝐺𝐻_ = +0.77 (p=1.49x10^-13^, Figure 4A). In contrast, we find a weaker negative correlation between −Δ𝐺°_𝐻20_values and *J(seq)* values, with a Pearson correlation coefficient of 𝜌_𝐺𝐽_ = −0.42 (p=7.5x10^-4^, Figure 4B). Although this negative correlation may be taken as evidence that positively covariant residues are destabilizing, it may alternatively reflect an underlying negative correlation between *H(seq)* and *J(seq)* values (Figure 4C). To isolate the correlation of *H(seq)* and *J(seq)* to stability from the indirect effects of correlation between *H(seq)* and *J(seq)*, we calculated partial correlation coefficients between *H(seq)* and −Δ𝐺°_𝐻20_ (𝜌_𝐺𝐻•𝐽_, equation 17) and between *J(seq)* and −Δ𝐺°_𝐻20_ (𝜌_𝐺𝐽•𝐻_, see Methods). The value of 𝜌_𝐺𝐻•𝐽_ is nearly the same as 𝜌_𝐺𝐻_ (0.73 versus 0.77), whereas the value of 𝜌_𝐺𝐽•𝐻_ is significantly smaller than 𝜌_𝐺𝐽_ (+0.15 versus -0.42). This indicates that *H(seq)* is the main determinant of stability for Potts designed HD sequences, whereas the correlation to *J(seq)* is an indirect effect of *H(seq)* on *J(seq)*.

**Figure 4.**
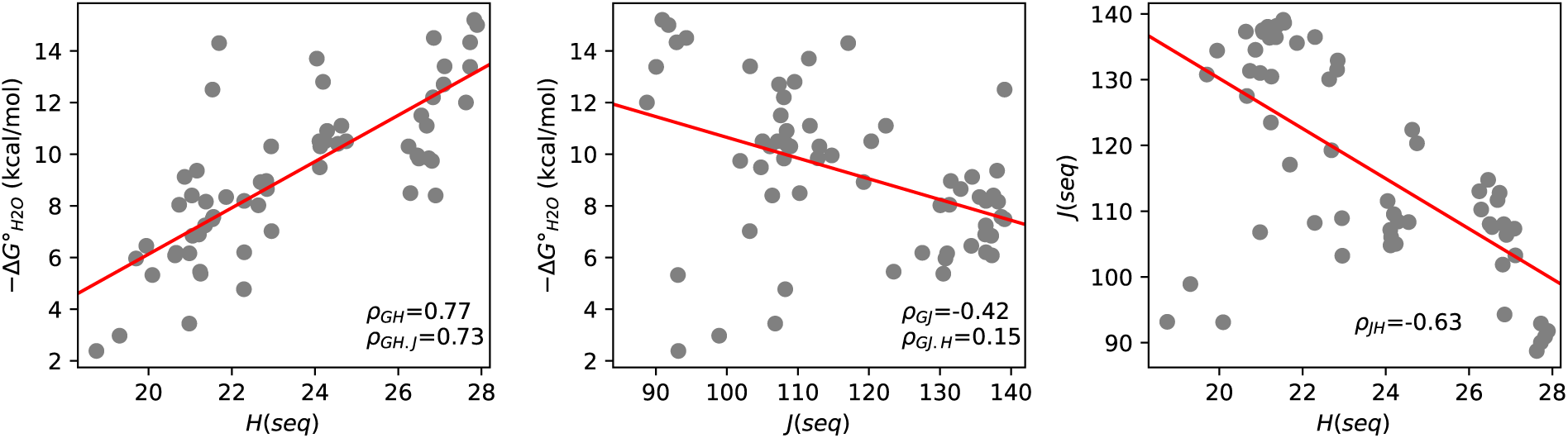
Correlation of stabilities of Potts-designed HD sequences with single-site and pair coupling scores. Stabilities (represented by negative ΔG°_H2O_ values) show a strong positive correlation (pearson correlation coefficient given as *ρ*_GH_, p=1.49x10^-13^) with *H(seq)* values (A), but a weaker negative correlation with *J(seq)* (B, p=7.5x10^-4^). Partial correlation coefficients (*ρ*_GH•J_, *ρ*_GJ•H_) indicate that the correlation between ΔG and *J(seq)* is indirect, resulting from a negative correlation between *J(seq)* and *H(seq)* (C). Sequences in this analysis include all Potts designs with all combinations of λ and X_ID_, as well as extant sequences (Tables S1A, S1B, and S3); values of *H(seq)* and *J(seq)* were all calculated using *h_i_(a)* and *j_i,k_(a,b)* coefficients generated using λ_h_=λ_j_=0.01, X_ID_=0.8.

### Stabilities of Potts-designed adenylate kinase sequences

To test whether our findings from the homeodomain family that the *h_i_(a)* coefficients are the primary determinants of protein stability are general, we performed Potts analysis on a second unrelated protein family, the enzyme adenylate kinase (AK). Because proteins in the AK family are significantly longer than proteins of the HD family (214 versus 57 residues in the MSA), there are many more pairs of residues in AK (22,791 versus 1,596); thus, we used the more aggressive regulation parameters (λ_h_=λ_j_=0.1) along with a sequence reweighting of X_ID_ = 0.8 for estimation of Potts coefficients. We used the resulting Potts coefficients in the *ɛ*(𝑠𝑒𝑞)_𝐻_, *ɛ*(𝑠𝑒𝑞)_𝐻𝐽_, and *ɛ*(𝑠𝑒𝑞)_𝐼_ energy functions (equations 9, 6, and 13) to optimize AK sequences with the Monte Carlo search procedure (Table S7), and expressed and purified the lowest energy sequence from each optimization.

As with most of the designed HD proteins, all three designed AK proteins expressed, were soluble, and have far-UV CD spectra consistent with the secondary structure of AK (Fig 5A). The stabilities of the designed AK sequences show the same trends as the stabilities of the designed HD sequences. Of the three AK proteins, the HJ-optimized sequence has the lowest stability, with a folding free energy of -11.2 kcal mol^-1^ and a C_m_ of 3.11 M GdnHCl (Fig 5B, Table S8). The H-optimized AK sequence significantly more stable than the HJ-optimized sequence, with an apparent folding free energy of -17.9 kcal mol^-1^ and a C_m_ of 5.50 M GdnHCl. Although the consensus AK protein generated from *ɛ*(𝑠𝑒𝑞)_𝐼_ optimization has the same fitted folding free energy as the H-optimized sequence (-18.1 and -17.9 kcal mol^-1^; Fig 5B, Table S8), the m-value for the H-optimized sequence is lower than that of the consensus protein (3.25 versus 5.67 kcal mol^-1^ M^-1^), indicating that folding of the H-optimized (and HJ-optimized) AK sequence may be multi-state. Thus, fitted folding free energies may not provide accurate representations of stability. As an alternative measure of stability, the C_m_ of the H-optimized AK sequence is significantly greater than that of the consensus protein (5.50 versus 3.87 M GdnHCl; Fig 5B, Table S8) demonstrating that the H-optimized sequence is significantly more resistant to GdnHCl denaturation than the consensus sequence. As with the Potts-designed HD sequences, the results with AK indicate that stability is maximized by optimizing *H(seq)*, exceeding the stability obtained either by optimizing *J(seq)* along with *H(seq)* or by consensus design.

**Figure 5.**
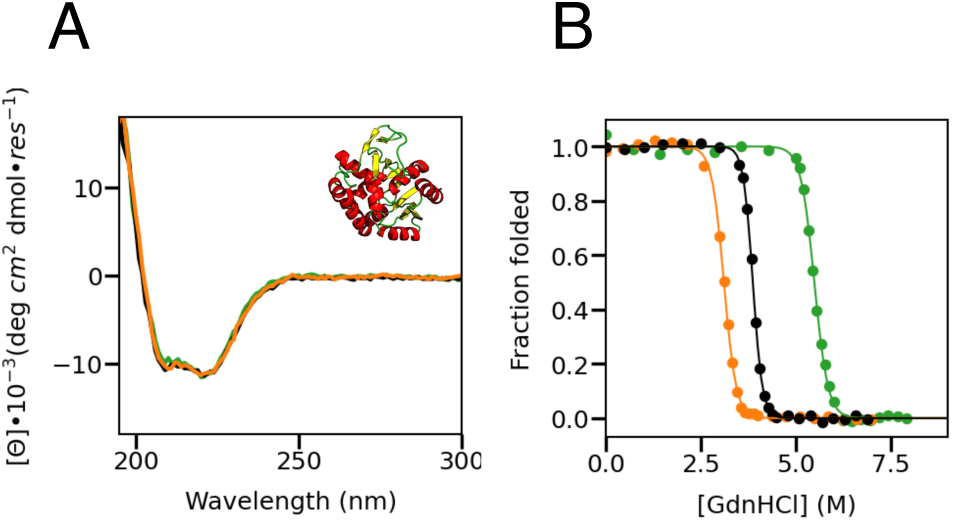
Structure and stability of Potts-designed adenylate kinases. (A) Far-UV CD spectra of lowest-energy AK sequences for H-(green), HJ-(orange), and I-optimization (black). Inset is a representative structure of *E. coli* AK (PDB: 1AKE). (B) GdnHCl-induced unfolding transitions of designed AK sequences (colors as in A). All transitions were collected at 20 °C. Solid lines represent the fit of a two-state unfolding model.

### Functional properties of Potts-designed proteins

Our results for the HD and AK families suggest that, on the whole, positively-covariant residue pairs do not increase protein stability, and to the extent that they conflict with single-site pair preferences, they may decrease stability. Since protein sequence alignments show clear statistical covariation patterns, it seems that covariance must contribute to some other aspect of protein fitness. We thus sought to determine how optimizing covariance (as opposed to pairwise conservation) impacts biological function.

The main function of homeodomains is to site-specifically bind to duplex DNA sequences to direct transcriptional activation.^22^ We thus determined DNA-binding affinities for H- and HJ-optimized sequences. Based on conserved sequence features, both proteins are predicted to bind to the 5’-TAATTA-3’ binding site typical for many HD families.^23^ Whereas all of these proteins retain the ability to bind DNA, H-optimized sequences bind with significantly higher affinity (K_d_ = 8.9 nM) than the HJ-optimized sequences (K_d_ = 343 nM, Figure 6, Table S11). This tighter binding affinity is achieved through a more favorable enthalpy of binding and is slightly offset by a less favorable entropy of binding. The H-optimized HD binds with similar binding affinity as a consensus HD we characterized in a previous study (K_d_ = 8.1 nM, Table S9), and is similar to the binding affinity of Ind1 (4.2 nM), whereas both the H- and HJ-optimized proteins bind with higher affinity than the extant *D. melanogaster* Engrailed HD (K_d_ = 787 nM, Table S9).^15^

**Figure 6.**
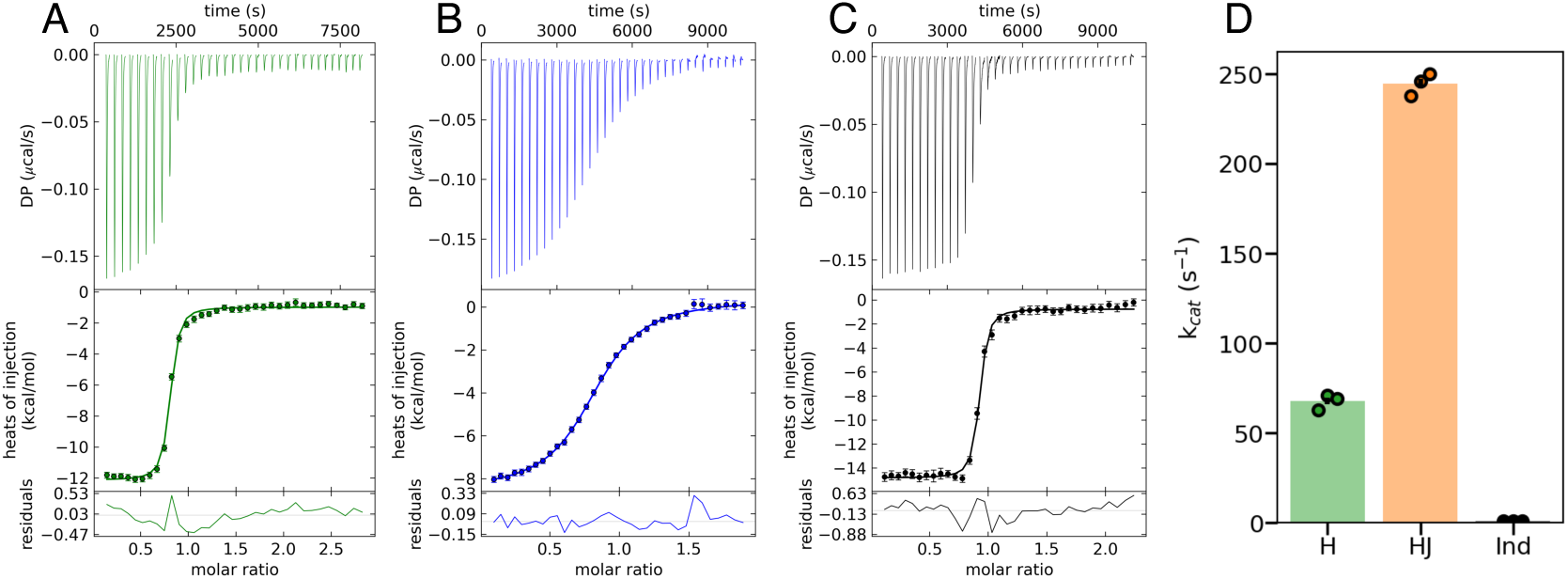
Functional analysis of Potts-designed proteins. (A-C) Isothermal titration calorimetry for designed HDs binding to DNA. HD sequences were generated using Potts coefficients λ_h_=λ_j_=0.01, X_ID_ = 0.8. Differential power (DP, top) and integrated heat peaks (middle) for titration of (A) H1, (B) HJA1, and (C) Ind1 into DNA at 20°C. Solid lines represent fits using a single-site binding model. (D) AK steady-state turnover numbers. Each point is the turnover number determined from a single replicate. Bars extend to the mean of the three replicates. AK activity measurements were carried out at 25 °C.

AK enzymes catalyze the reversible conversion of ATP and AMP to two molecules of ADP. We determined the turnover number in the direction of ADP formation under steady-state conditions for all three designed AK proteins. Although all designed AKs retain measurable catalytic activities for the phosphotransfer reaction (Figure S10), there are significant activity differences among the three proteins. Both the H and the HJ-optimized AK proteins have significantly higher turnover numbers (244 and 68 sec^-1^) than consensus AK (1.01 sec^-1^; Figure 6B, Table S10); both of these values are within the range seen for extant AK enzymes under similar conditions (Table S10).

## Discussion

The above results indicate that to the extent that Potts design increases protein stability, it does so not by including positively covariant residue pairs, but by including single-site biases. Consensus design, which has been shown to generate stable proteins^24,25^ also generates sequences that capture single-site biases; however, because positively covariant residues tend to occur frequently, consensus design is expected to inadvertently include positively covariant residues. The Potts model provides a way to separate single-site biases from and pair correlations. Our finding that sequences generated by optimizing *H(seq)* are more stable than those generated with partial or full *J(seq)* optimization is consistent with the interpretation that single-site biases rather than pairwise coupling stabilize proteins.

The observation that pairwise couplings do not contribute to protein stability is surprising. Inferred couplings from the Potts model have been shown to correlate with structural contacts^20,26^ and have been important for recent advances in protein structure prediction. Since protein structures are generally considered to be free energy minima^27^, pairwise couplings might be expected to help define such minima. Moreover, analysis of deep mutational screens have showed a positive correlation between various aspects of protein fitness (including stability) and Potts energy scores^28–30^, and design using Potts energy scores have been shown to generate stable folded proteins.^31^ One explanation for this apparent discrepancy is that these studies have optimized both *h_i_(a)* and *j_i,k_(a,b)* scores, and have compared these proteins to extant proteins. Indeed, when we compare stabilities of extant proteins with HJ-optimized sequences, in which *h_i_(a)* and *j_i,k_(a,b)* scores are equally weighted, the stabilities of the HJ-optimized sequences are more than the extant sequences. This stabilization is likely the result of the modest increases in *H(seq)* values (Figure 2) that result from HJ-optimization. Although our designed sequences that maximize couplings are more stable than extant sequences, they are considerably less stable than those in which *H(seq)* values are maximized at the expense of *J(seq)* values.

Previous work by Ranganathan and coworkers indicate that including residue coupling is necessary to generate artificial sequences that properly fold.^2^ They found that artificial WW domain proteins designed to preserve only the single-site residue frequencies were not folded (0 out of 43 designs), whereas designs that preserved covariance were more often folded (12 out of 43 designs). Although these results seem at odds with our findings, the Ranganathan study designed sequences to preserve the average biases of extant proteins, whereas our strategy maximizes these biases.

Our findings single-site *h_i_(a)* values determine thermodynamic stability is consistent with a recent high-throughput study from Lehner and coworkers that showed that protein abundance in a yeast expression system could be well-explained by a simple thermodynamic model including only single-site biases^32^. However, in that study, including pairwise couplings improved the accuracy of the thermodynamic model, consistent with previous deep-mutational studies^28–30^ but at least superficially at odds with the results here. One possible explanation for this discrepancy is that the improved accuracy from including pairwise terms in the Lehner study need not reflect a favorable contribution of pairwise correlations to stability, but may provide a more accurate determination of the single-site biases.

While the residue coupling information does not appear to contribute to protein stability, it may be important for protein function. For the HD and AK families, sequences designed optimizing single-site information alone as well as optimizing single-site and coupling information maintained expected biological activities. For the HD family, the protein optimizing *J(seq)* has a higher DNA binding affinity than the extant Drosophila Engrailed HD, although not to the same extent as the protein optimizing *H(seq)*. For AK activity, the protein optimizing *J(seq)* increased k_cat_ by 240-fold compared to the consensus AK protein. It is rather surprising that the protein optimizing *H(seq)* also increased k_cat_ significantly compared to the consensus protein, although not to the same level as for the HJ-optimized protein. Determining whether *J(seq)* scores determine enzyme activity will require a more systematic analysis of Potts constructs in the AK and other enzyme families, but our findings here are consistent with studies linking Potts scores to enzyme activity^28^. We note that our results with AK activity are not consistent with the stability-activity tradeoff relationship that has been described for other enzymes.

The observation that stability can be increased by maximizing single-site Potts energies has implications for protein design. Optimizing stability using single-site energies may provide a route to further enhance stabilities beyond the stabilization typically afforded by consensus design^25^, and may provide a route to stabilization in what seems to be the minority of cases where consensus sequences do not increase stability^33^. One limitation of consensus design is that it appears to generate enzymes with decreased catalytic proficiencies.^24^ To the extent that H-optimized sequences retain (or have enhanced) rates of catalysis, as we have found for H-optimized AK, the use of Potts-based H-optimization may provide a general route to enzymes that are both highly stable and highly active.

## Materials and Methods

### Obtaining multiple sequence alignments

For the HD family, we obtained the “Full” alignment from Pfam (PF00046, accessed on 10/23/18).^34^ Positions with gap frequencies greater than 50% were removed, resulting in a sequence length of L=57 for all sequences. Resulting sequences that contained greater than 50% gap characters were removed from the alignment, as were identical sequences. The resulting in an alignment contained m=19,221 sequences.

For the AK family, we obtained sequences from the InterPro database (IPR007862, accessed on 5/13/19).^35^ Sequences containing nonstandard amino acids and sequences shorter than 172 residues or longer than 258 residues were removed from the sequence set. Sequences were then aligned using MAFFT.^36^ Positions with gap frequencies greater than 50% were removed, resulting in a sequence length of L=214 for all sequences. Sequences that contained greater than 10% gap characters were removed from the alignment, as were identical sequences. The resulting alignment contained m=14,090 sequences.

### Inference of single-site and coupling energy coefficients using the Potts model

Single-site and pairwise coupling energies were inferred from MSAs using a Potts-like formalism.^5^ In this formalism, sequence probabilities are assumed be governed by an equilibrium

Boltzmann distribution:

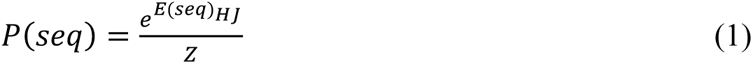

where Z is a normalization constant (similar to a partition function in statistical mechanics) such that probabilities of all sequences sum to one. *E(seq)_HJ_* is the energy of a sequence computed from single-site and pairwise coupling energies:

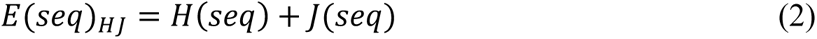

where *H(seq)* is the total single-site energy and *J(seq)* is the total pairwise coupling energy for a given protein sequence:

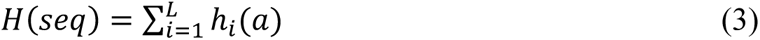

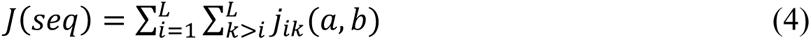

Note that in this formalism, favorable *h_i_(a)* and *j_ik_(a,b)* terms have large positive values; thus the exponent in the Boltzmann expression (1) lacks a negative sign.

To infer the 𝐿 × 21 (for the 20 amino acids plus a gap) single-site energy coefficients and the 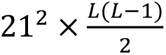 coupling energy coefficients, we used the pseudolikelihood optimization procedure of Aurell and coworkers.^37,38^ Due to the finite sampling of sequences in the MSA combined with the large number of *j_ik_(a,b)* coefficients, the Potts model is prone to overfitting. To mitigate overfitting, the objective function used to infer *h_i_(a)* and *j_ik_(a,b)* coefficients contains an ℓ_2_ regularization penalty term given as:

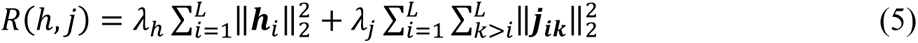

where 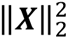 is the squared ℓ_2_-norm of coefficient matrix **X**, and λ_h_ and λ_j_ are parameters tuning the magnitude of the regularization for the single-site and coupling coefficients respectively. In separate fits of the HD MSA, we applied values of λ_h_=λ_j_=0.1 and λ_h_=λ_j_=0.01 to explore the effects of the magnitude of regularization on sequences generated from the Potts formalism and their properties. Because AK is significantly longer than HD and thus has many more residue pairs, we applied the more aggressive regularization (λ_h_=λ_j_=0.1) for Potts analysis of AK.

To decrease the effect of phylogenetic biases on the sequence composition of the MSAs, the contribution of each sequence to the model objective function was weighted based on sequence identities. A threshold sequence identity (X_ID_) was chosen to identify sequences with high similarities. For a sequence *b* in the MSA, we determined the number of sequences in the MSA, *m_b_*, with identities to sequence *b* greater than the threshold X_ID_, and weighted the contribution of sequence *b* in the pseudolikelihood function by 𝑤_𝑏_ = 1⁄𝑚_𝑏_. Thus, a sequence with high similarity to many other sequences in the MSA (average pairwise identity above X_ID_) is given low weight. For the HD MSA, we used sequence identity thresholds of X_ID_ = 0.2 and X_ID_ = 0.8 to examine the effects from sequence weighting on the model fit. For the AK MSA, we used a value of X_ID_ = 0.8.

### Sequence design using single-site and coupling energy energies

To design sequences using the single-site and pairwise coupling energies determined from the Potts model, we performed simulated annealing Monte Carlo simulations similar to those described by Best and coworkers.^9^ We defined a set of Monte Carlo energy functions *ɛ*(𝑠𝑒𝑞) that combine the single-site and pairwise coupling energies in different proportions:

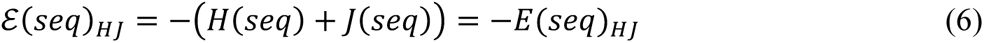

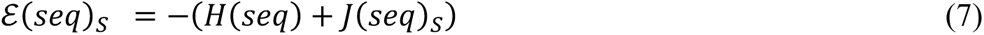

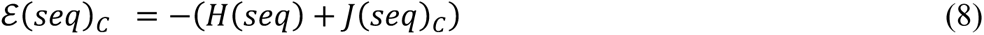

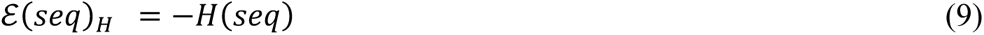

The sign convention in equations 6-9 inverts the Potts energy scores so that the most stabilizing scores lead to low values of *ɛ*(𝑠𝑒𝑞) in the Monte Carlo sequence optimization. The *ɛ*(𝑠𝑒𝑞)_𝐻𝐽_ energy function includes all single-site and pairwise coupling energies and is equivalent to the Potts energy function used for maximum likelihood estimates of *h_i_(a)* and *j_ik_(a,b)* terms (equation 1). The *ɛ*(𝑠𝑒𝑞)𝑆 energy function is analogous to *ɛ*(𝑠𝑒𝑞)_𝐻𝐽_, but it only includes the strongest pairwise coupling energies. Strongly coupled pairs were identified by calculating the average product-corrected Frobenius norms of the 21x21 coupling coefficient matrices (‖𝑗_𝑖𝑘_‖_𝐴𝑃𝐶_) for all residue pairs, and selecting only the pairs with norms in the top 10 percent. Similarly, the *ɛ*(𝑠𝑒𝑞)𝐶 energy function only includes only *j_ik_(ab)* values for residues that are close to one another in the homeodomain structure. Close residue pairs were identified by determining the distances between all pairs of Cβ atoms (Cα atoms for glycines) in the *D. melanogaster* Engrailed HD structure (PDB: 2JWT) and selecting only the pairs with distances in the top 10 percent. These two *j_i,k_(a,b)* filters were implemented using a Heaviside step function 𝜃:

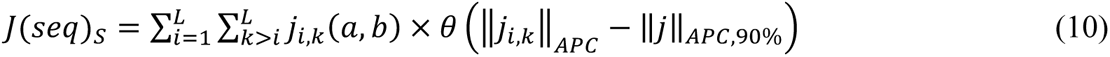

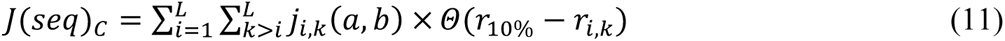

Where ‖*j_i,k_*‖_𝐴𝑃𝐶_ is the average product corrected Frobenius norm of the coupling matrix between the *i*^th^ and *k*^th^ positions, ‖𝑗‖_𝐴𝑃𝐶,90%_ is the 90^th^ percentile average product corrected Frobenius norm, *r_i,k_* is the Cβ-Cβ distance of the of the *i*^th^ and *k*^th^ position pair, and r_10%_ is the Cβ-Cβ distance at the 10^th^ percentile rank. The *ɛ*(𝑠𝑒𝑞)𝐻 energy function uses only the *h_i_(a)* coefficients determined from the maximum likelihood estimation of Potts coefficients (*h_i_(a)* and *j_i,k_(a,b)* terms), and can be considered to be an energy function dependent only on single-site information that is free from inadvertant statistical biases produced by pair correlations.

In addition to generating sequences using various combinations of the Potts coefficients *h_i_(a)* and *j_i,k_(a,b)*, we generated sequences using coefficients from an independent model in which pair correlations are ignored during pseudolikelihood optimization. In this optimization, sequence probabilities are assumed to be governed by a Boltzmann model analogous to equation 1:

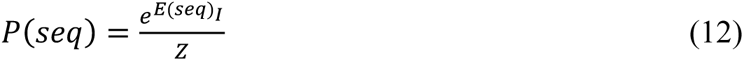

where

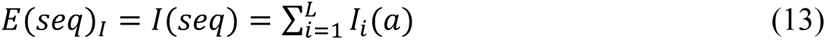

The 𝐼_𝑖_(𝑎) coefficients from the independent model are analogous to the ℎ_𝑖_(𝑎) of the Potts model in that they are single-site coefficients. However, these two sets of coefficients differ in that the ℎ_𝑖_(𝑎) coefficients were inferred in a model that explicitly analyzed biases from pair correlations and separated these biases into the *j_i,k_(a,b)* terms, whereas pair biases are ignored in the independent model, and end up contributing to the 𝐼_𝑖_(𝑎) coefficients.

Sequences were generated from the independent model using the Monte Carlo search procedure described below with the energy function

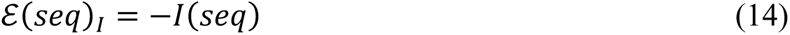

*I-*optimized sequences are closely analogous to consensus sequences, where residue selection at each position ignores the surrounding sequence context, and is inadvertently influenced by biases from pair correlations.

Using the energy functions defined above (equations 6-9, 14), Monte Carlo sequence generation was initiated from a random sequence generated by choosing (with uniform probabilities) a non-gap residue at each position from the set of residues found at the same aligned position in the MSA. At each Monte Carlo step, one residue is randomly chosen and substituted with a different non-gap residue found at the same position in the MSA, resulting in a substituted sequence. The residue substitution is accepted with a probability

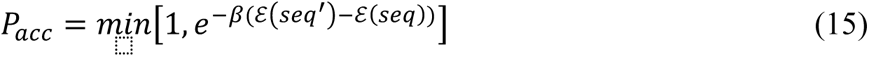

where *ɛ*(𝑠𝑒𝑞) and *ɛ*(𝑠𝑒𝑞′) are energies of starting and substituted sequences. Accordingly, a substitution that lowers the energy will always be accepted, whereas a substitution that raises the sequence energy will be accepted with a Boltzmann-weighted probability based on the difference in energies the two sequences. *β* is a factor that scales the energy change associated with sequence substitution, analogous to an inverse temperature. Each simulation begins at a *β* value of 0.1, where unfavorable substitutions are accepted with high probabilities. Each simulation runs for 100,000 Monte Carlo steps, and *β* is increased (temperature decreases) every 5,000 steps such that probability of accepting unfavorable substitutions decreases. The increment in *β* is chosen to ensure convergence of the simulation to low-energy sequences by the end of the simulation. For each energy function, we run 1,000 independent simulations starting from different randomly generated sequences, producing 1,000 sequences optimized by the given energy function.

### Gene synthesis, protein expression, and preparation

Gene sequences encoding proteins studied were synthesized by Twist Bioscience and cloned into pET28 expression vectors between the *NcoI* and *XhoI* restriction sites. HD constructs included sequences encoding an N-terminal Met-Gly-Ser sequence for translation initiation and cloning, as well as a C-terminal His_6_-tag for purification. AK constructs included sequences encoding a Gly-Ser-Trp sequence inserted after the N-terminal Met for cloning and quantification, as well as a C-terminal His_6_-tag for purification.

*E. coli* BL21(DE3) cells containing plasmids for HD and AK expression were grown at 37 °C to an OD_600_ of 0.6-0.8 and induced with IPTG at a final concentration of 1 mM. HDs were expressed overnight at 20 °C and AKs were expressed for 4-6 hours at 37 °C. Cell pellets were harvested by centrifugation and resuspended in buffer containing 50 mM Tris (pH 8.0) for HDs or 50 mM Tris (pH 8.0) and 1 mM TCEP for AKs. Buffers were supplemented with a cocktail of protease inhibitors (Roche cOmplete EDTA-free) to inhibit protein degradation. Cells were lysed by sonication and the cell lysate was clarified by centrifugation. The supernatant was collected, supplemented with MgCl_2_ and CaCl_2_ to a final concentration of 2 mM for each, and incubated with 1 mg DNaseI and 250 units of Benzonase nuclease for 1-3 hours at room temperature. Proteins were then purified by Ni-NTA and cation exchange chromatographies. Purified HDs were dialyzed into buffer containing 25 mM NaPO_4_ (pH 7.0) and 150 mM NaCl, and purified AKs were dialyzed into 25 mM Tris (pH 8.0), 50 mM NaCl, and 1 mM TCEP.

Two of the AK proteins (the single-site optimized protein and the consensus protein) were found to co-purify with endogenous substrate after undergoing the above protocol. These proteins were loaded back onto the NiNTA column, washed with buffer containing 25 mM Tris (pH 8.0), 50 mM NaCl, 1 mM TCEP, and 6-8 M guanidine hydrochloride (GdnHCl) to unfold the proteins, thereby dissociating and removing substrate, washed with buffer containing 25 mM Tris (pH 8.0), 50 mM NaCl and 1 mM TCEP to refold the proteins on the Ni-NTA column, then eluted off the column under native conditions. Purified proteins were then dialyzed back into buffer containing 25 mM Tris (pH 8.0), 50 mM NaCl, and 1 mM TCEP. All proteins were flash frozen in liquid nitrogen and stored at -80 °C.

### Circular dichroism spectroscopy

CD experiments were performed using Aviv spectropolarimeters. Far-UV CD spectra were collected at 25 °C using an 0.1-cm cuvette. For HDs, samples were prepared with protein concentrations of 10-15 uM in buffer containing NaPO_4_ (pH 7.0) and 150 mM NaCl. For AKs, samples were prepared with protein concentrations of 3 uM in buffer containing 25 mM Tris (pH 8.0), 50 mM NaCl, and 0.5 mM TCEP.

Equilibrium GdnHCl-induced unfolding was monitored by CD at 222 nm. For all AKs, protein samples at varying concentrations of GdnHCl were incubated for two days at room temperature and CD was read for each sample. Protein concentrations ranged from 1-3 uM and samples were read at 20 °C. For HDs, a Hamilton automated titrator was used for GdnHCl titrations. Samples were allowed to equilibrate for five minutes after each injection of titrant. Protein concentrations ranged from 6-12 uM and samples were read at 25 °C. Folding free energies in the absence of GdnHCl were determined using a two-state linear extrapolation model.^39^

For the most stable HD proteins, GdnHCl titrations at 25°C inadequately resolved the unfolded baseline, preventing an accurate determination of folding free energies. To better define unfolded baselines, we combined GdnHCl titrations at 25 °C with titrations at three elevated temperatures, and fit all four unfolding transitions to a global model to determine a folding free energy at 25 °C. In the global model, each unfolding transition was fit with a local native baseline, a free energy in the absence of denaturant, and an m-value parameters, but all four unfolding transitions were fit to a common unfolded baseplane (s_unfolded_):

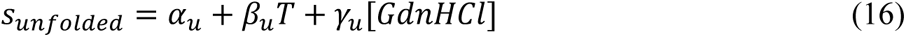

where α*_u_* is the intercept of the unfolded baseplane at zero Kelvin and in the absence of denaturant, β*_u_* is slope of the baseplane with respect to temperature, and γ_u_ is the slope of the baseplane with respect to GdnHCl. To minimize melt-to-melt variability, all four denaturation experiments were performed successively on the same spectrophotometer with the same protein sample and titrant stocks.

### Correlation analysis between HD stability, *H(seq)*, and *J(seq)*

As shown in Figure 4, the correlations between −Δ𝐺°_𝐻20_ and the *H(seq)* and *J(seq)* scores may be the indirect result of correlation between *H(seq)* and *J(seq)*. To isolate the effects of *H(seq)* and *J(seq)* on −Δ𝐺°_𝐻20_ from the potentially confounding effects of the other variable, we calculated partial correlation coefficients, 𝜌_𝐺𝐻•𝐽_ and 𝜌_𝐺𝐽•𝐻_. Numerically, 𝜌_𝐺𝐽•𝐻_ is the correlation coefficient between the residuals from linear regression of −Δ𝐺°_𝐻20_ on the *H(seq)* scores (𝑟_𝑖,𝐺𝐻_, Figure 4A) and the residuals from the regression of the *J(seq)* scores on the *H(seq)* scores (𝑟_𝑖,𝐽𝐻_, Figure 4C):

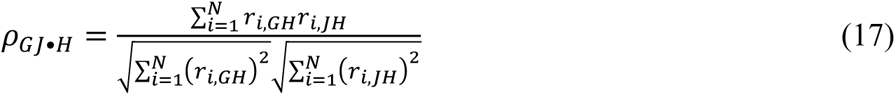

The residuals in linear regression represent the variation in the dependent variable that cannot be accounted for from variation of the independent variable. Thus, a large value of 𝜌_𝐺𝐽•𝐻_ indicates that the variations in −Δ𝐺°_𝐻20_ that are independent of the *H(seq)* scores (represented by the 𝑟_𝑖,𝐺𝐻_ values) are correlated with the variations in the *J(seq)* scores that are independent of the *H(seq)* scores (represented by 𝑟_𝑖,𝐽𝐻_ values). Conversely, a small value of 𝜌_𝐺𝐽•𝐻_ indicates that the variations in −Δ𝐺°_𝐻20_ that are independent of *H(seq)* are also independent of the variation in *J(seq)* values that are independent of *H(seq)*. That is, removing the correlation of *J(seq)* with *H(seq)* also removes the correlation of −Δ𝐺°_𝐻20_with *J(seq)*, implying that the apparent correlation of −Δ𝐺°_𝐻20_ with *J(seq)* is an indirect result of correlation of −Δ𝐺°_𝐻20_ with *H(seq)*.

### Isothermal titration calorimetry of HD DNA binding

The self-complementary DNA oligonucleotide 5’-CGACTAATTAGTCG-3’ was purchased from IDT. Oligonucleotides were resuspended in buffer containing 25 mM NaPO_4_ (pH 7.0) and 250 mM NaCl, and duplex DNA was formed by incubating the oligonucleotide at 95 °C for 5 minutes and slowly cooling to room temperature. Protein and DNA stocks were dialyzed against the same buffer, 25 mM NaPO_4_ (pH 7.0) and 250 mM NaCl. The concentrations of protein and DNA samples were determined after dialysis by UV absorbance. For all titrations, protein concentrations between 30 – 95 μM were injected into DNA samples at a concentrations 10 times lower than the protein (3 - 9 μM). All titrations were performed at 20 °C. Thermograms and binding isotherms were analyzed using the SEDPHAT software suite and fit to a single site binding model to determine DNA binding affinities.^40^

### AK steady-state enzyme kinetics

Steady-state kinetic rates under saturating substrate conditions were determined using a coupled spectroscopic assay in the direction of ADP formation.^41^ Reactions were carried out in 50 mM HEPES (pH 7.5), 100 mM NaCl, 20 mM MgCl_2_, 0. 5mM TCEP, 10 mM phosphoenolpyruvate, 0.1 mM NADH, 7.4 mM AMP, 9.2 mM ADP, 27-42 units/mL lactate dehydrogenase, and 18-30 units pyruvate kinase. All reactions were collected at 25 °C. Reactions were initiated by addition of AK enzyme to final concentrations of 0.9 nM for the single-site optimized AK, 0.4 nM for the single-site and coupling optimized AK, and 122 nM for the consensus AK.

## Acknowledgments and Funding Sources

We thank members of the Barrick lab for discussions of experimental design, data analysis, and interpretations presented here. This work was supported by NIH/NIGMS Grant 1R01 GM068462 to DB, an NIH Training grant T32 GM008403, and NIH/NIGMS fellowship F31 GM128295 to MS.

## Supplementary Figures and Tables

**Figure S1.**
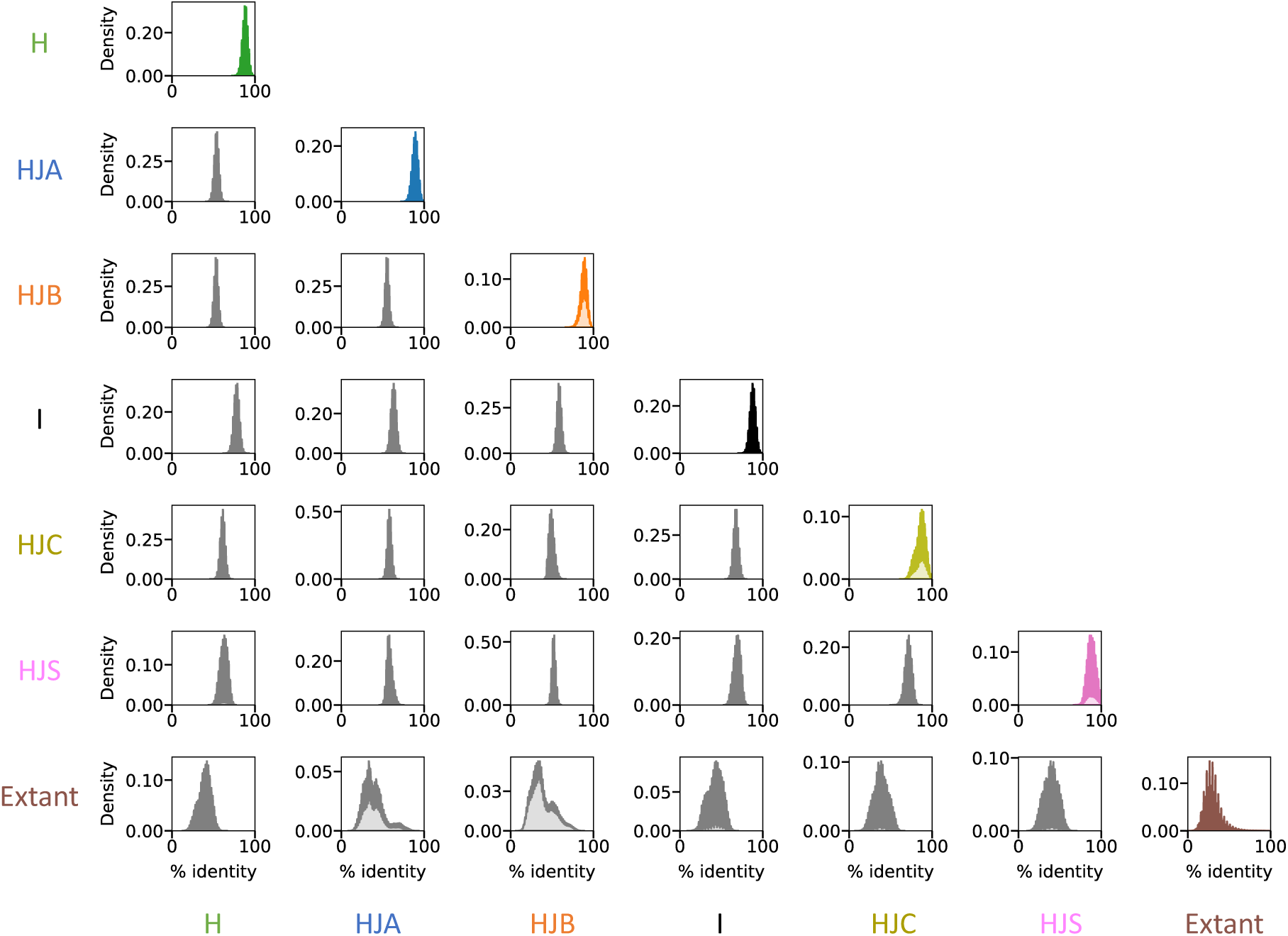
Identities among groups of designed and extant HD sequences. Plots on the diagonal show all pairwise sequence identities among 1000 sequences optimized with the given energy function or among extant sequences (bottom row). Plots in the lower triangle (grey) show pairwise sequence identities between different sequence groups. Sequences were generated with a sequence reweighting X_ID_=0.8 and regularization parameters λ_h_=λ_j_=0.01.

**Figure S2.**
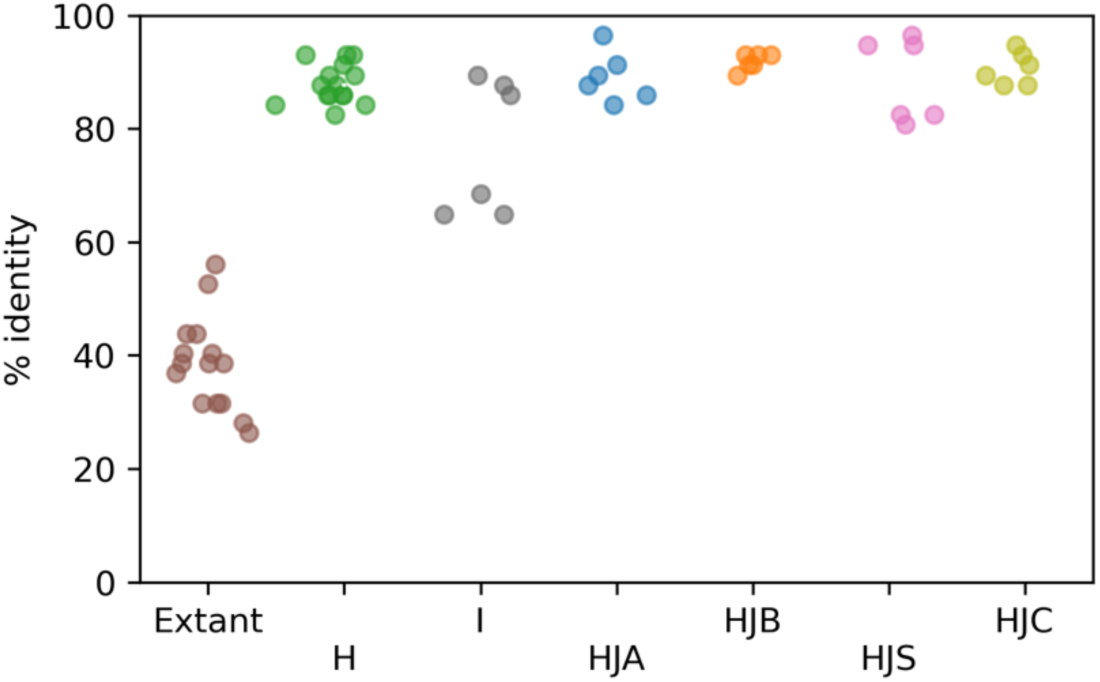
Sampled HD sequence identities. Pairwise sequence identities among the HD sequences selected for biochemical characterization, including sequences derived from the Potts energy functions (H, HJA HJB, HJS, and HJC), from the independent model (I) and extant sequences from the multiple sequence alignment. Sequences were generated with a sequence reweighting X_ID_=0.8 and regularization parameters λ_h_=λ_j_=0.01.

**Figure S3.**
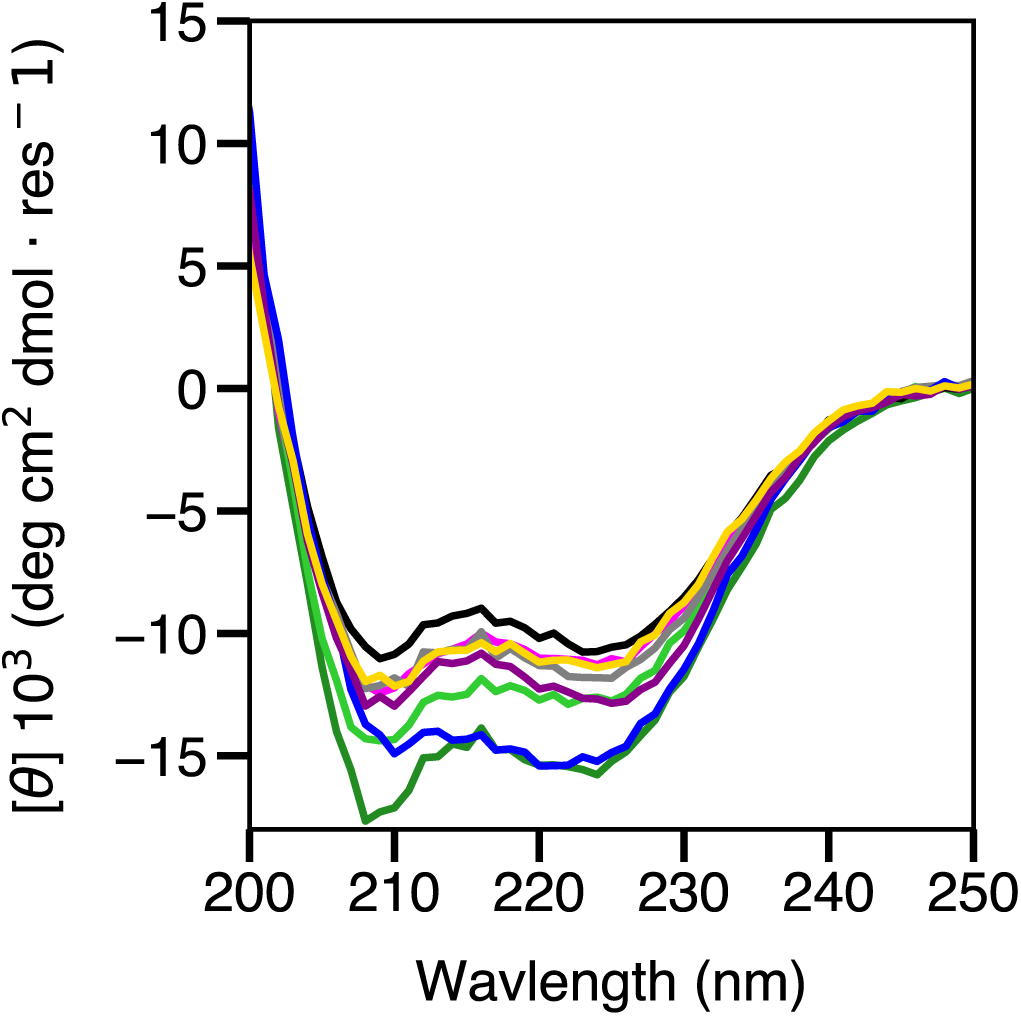
Circular dichroism spectra of Potts-designed HD proteins with sequence reweighting X_ID_=0.8, λ_h_=λ_j_=0.01. Spectra were collected at 25 °C for H1 (green), H4 (light green), Ind1 (black), Ind3 (gray), HJA1 (blue), HJs1 (magenta), HJS3 (light magenta), HJC1(yello) Potts-designed proteins.

**Figure S4.**
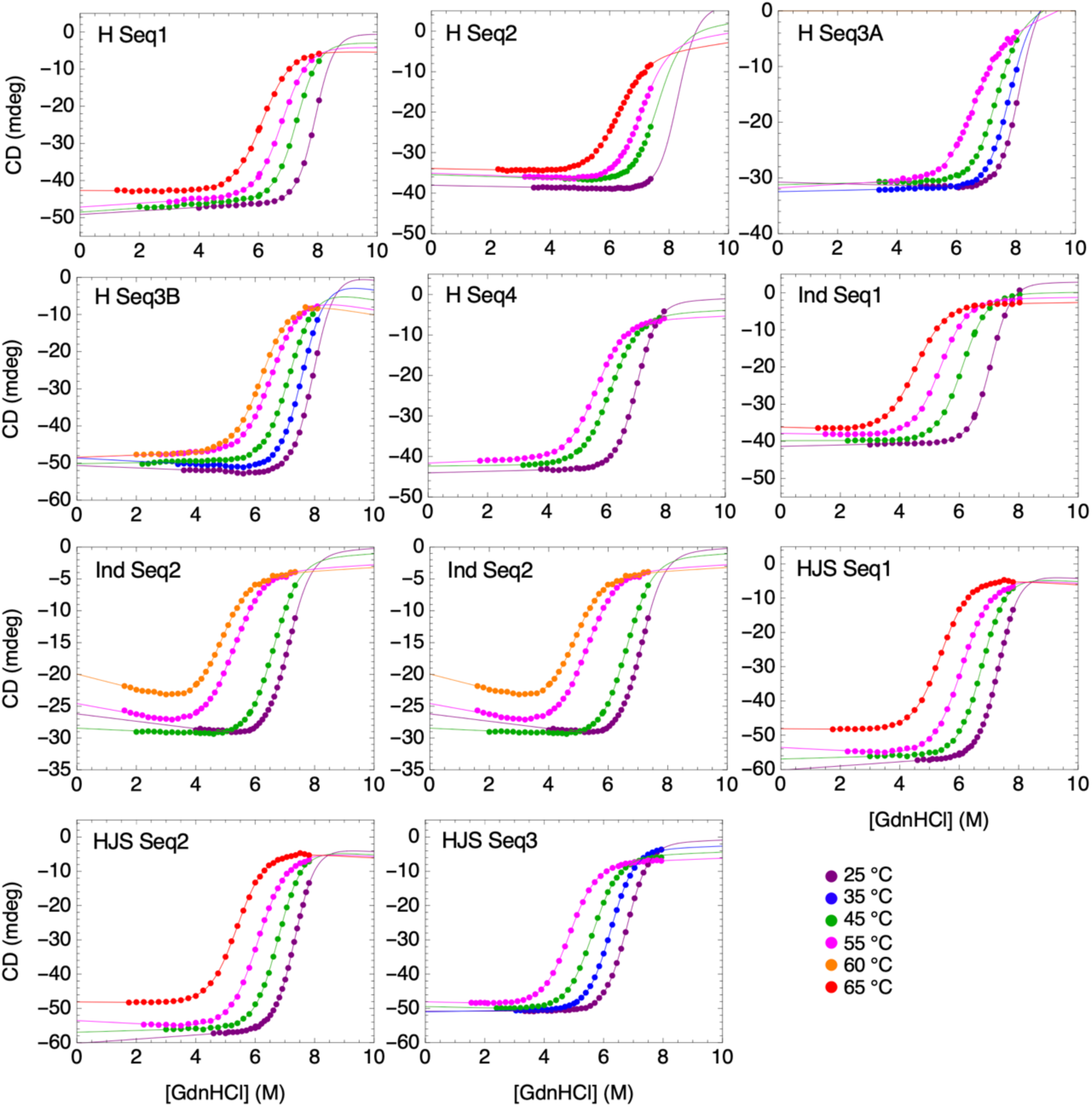
GdnHCl induced unfolding transitions at different temperatures for HDs with high stabilities. Protein identity and Potts model parameterization are indicated inset in each plot; temperatures are indicated in the legend, right. Solid lines are from fits of a global model to all four unfolding transitions using a common denatured baseplane. Each unfolding transition contains local native baseline parameters, local free energies, and local m-values. Sequences were generated using regularization coefficients λ_h_=λ_j_=0.01 and a sequence reweighting threshold of X_ID_=0.8.

**Figure S5.**
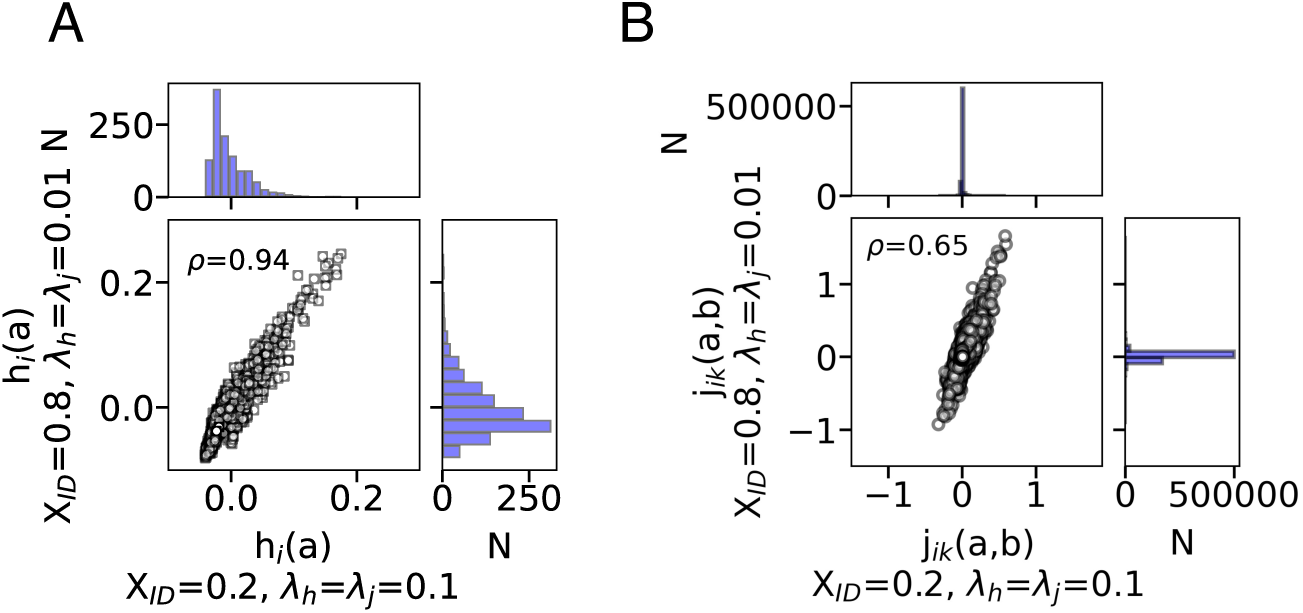
The effects of model parameterization on inferred Potts coefficients. Comparison of all single-site (A) and coupling (B) energy coefficients from fitting the Potts model with different values of sequence identity reweighting threshold X_ID_ and regularization coefficients λ_j_=λ_h_ to the same HD MSA. Pearson correlation coefficient (*ρ*) shown for each comparison is shown on each plot.

**Figure S6.**
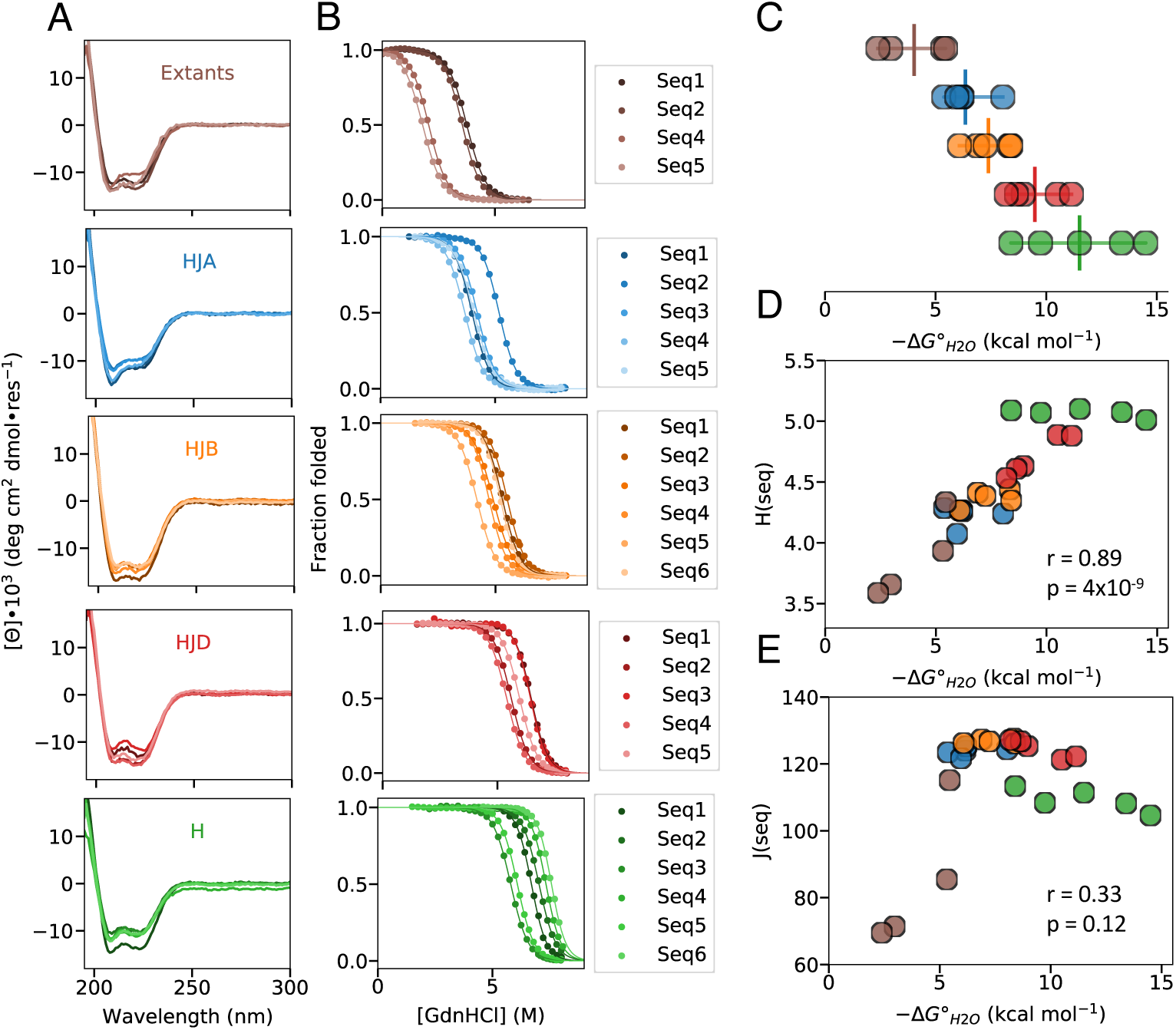
Stabilities of Potts-designed HDs with X_ID_=0.2, λ**_j_=**λ**_h_=0.1.** (A) Far-UV CD spectra of exatant and Potts-designed HDs. (B) Representative GdnHCl-induced unfolding transitions of HDs at 25 °C. Solid lines represent the fit of a two-state unfolding model; for sequences with incomplete high-temperature baselines, fits included additional GdnHCl transitions (Figure S7). (C) Folding free energies of HDs determined from the two-state unfolding analyses as in panel B. Vertical bar indicates the mean folding free energy of each distribution. (D and E) Correlation of folding free energies and sequence single-site energies (D) and sequence coupling energies (E).

**Figure S7.**
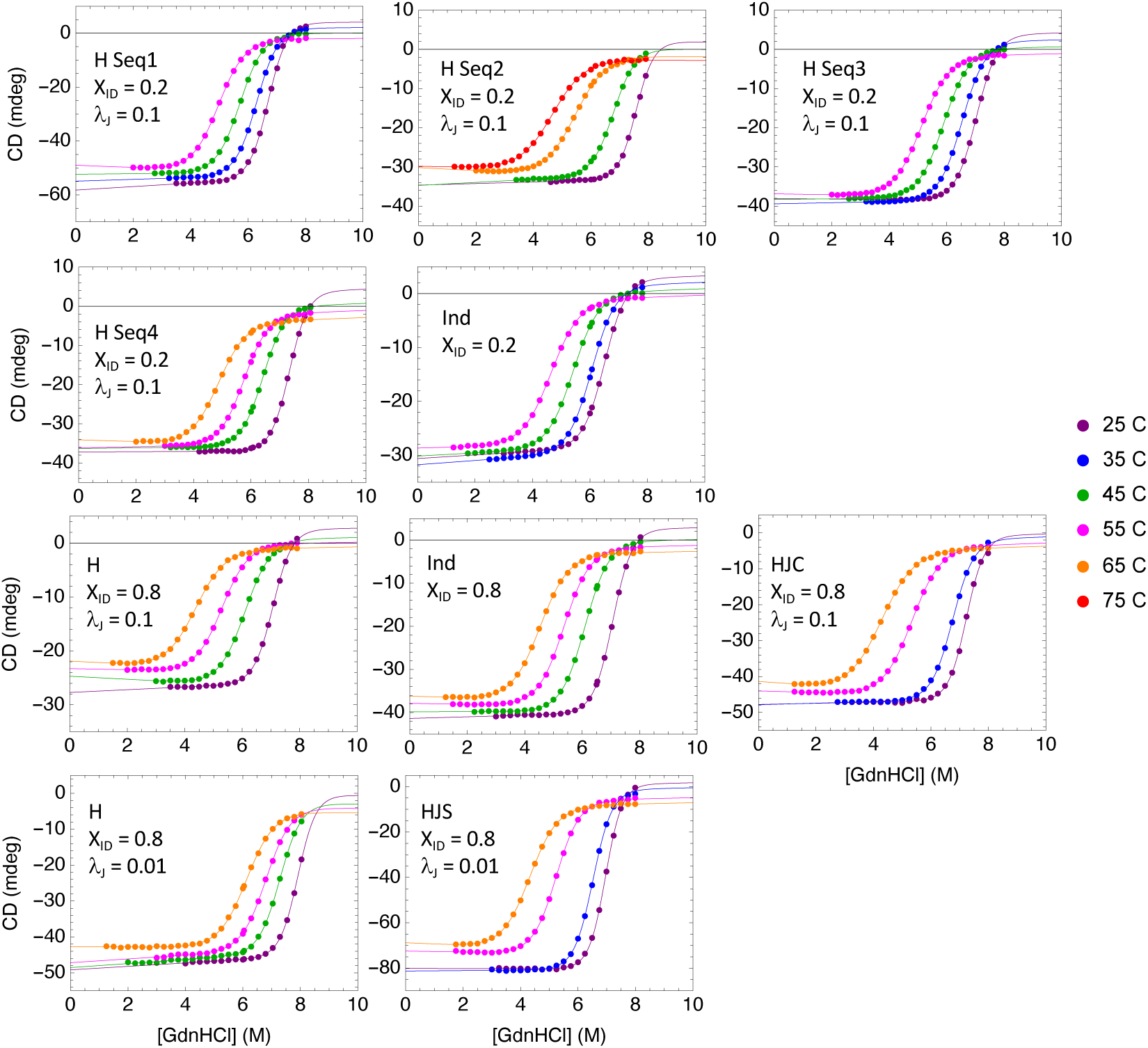
GdnHCl induced unfolding transitions at different temperatures for HDs with high stabilities. Protein identity and Potts model parameterization are indicated inset in each plot; temperatures are indicated on the legend, right. Solid lines are from fits of a global model to all four unfolding transitions using a common denatured baseplane. Each unfolding transition contains local native baseline parameters, local free energies, and local m-values. Sequences were generated using regularization coefficients λ_h_=λ_j_=0.1 and a sequence reweighting threshold of X_ID_=0.2.

**Figure S8.**
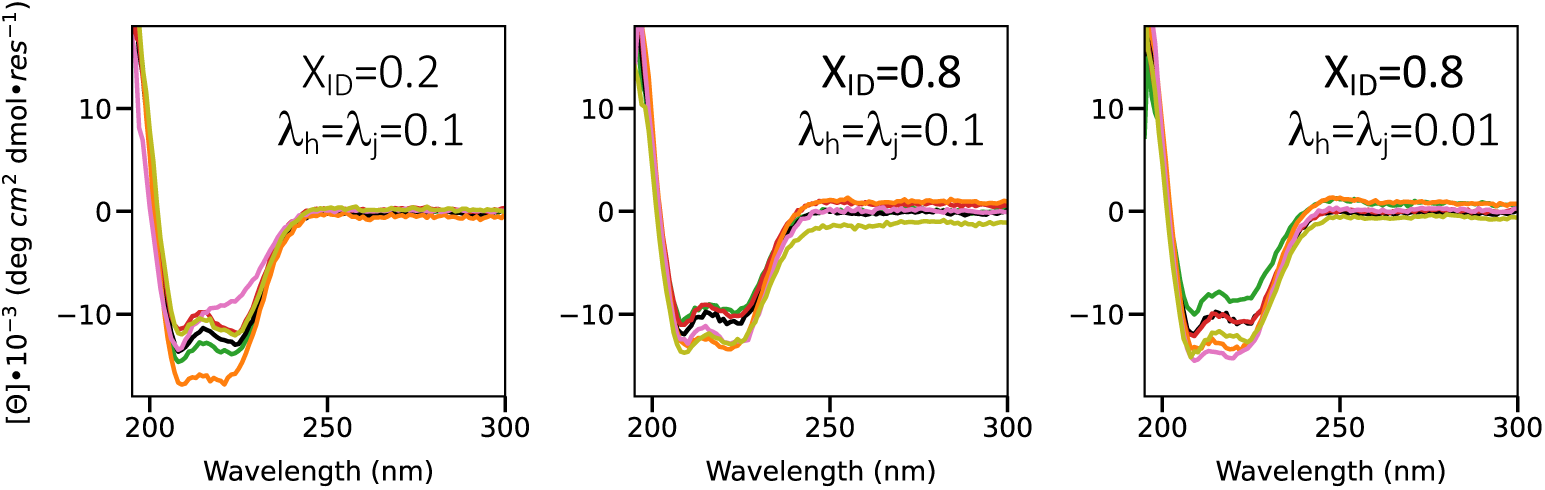
Far-UV CD spectra of Potts-designed HD proteins with different global Potts fitting parameters. Sequence identity reweighting thresholds X_ID_ and regularization coefficients (λ_j_=λ_h_) are given in each panel. Spectra were collected at 25 °C for H (green), HJ (orange), Ind (black), HJD (red), HJS (pink), HJC (gold) Potts-designed proteins.

**Figure S9.**
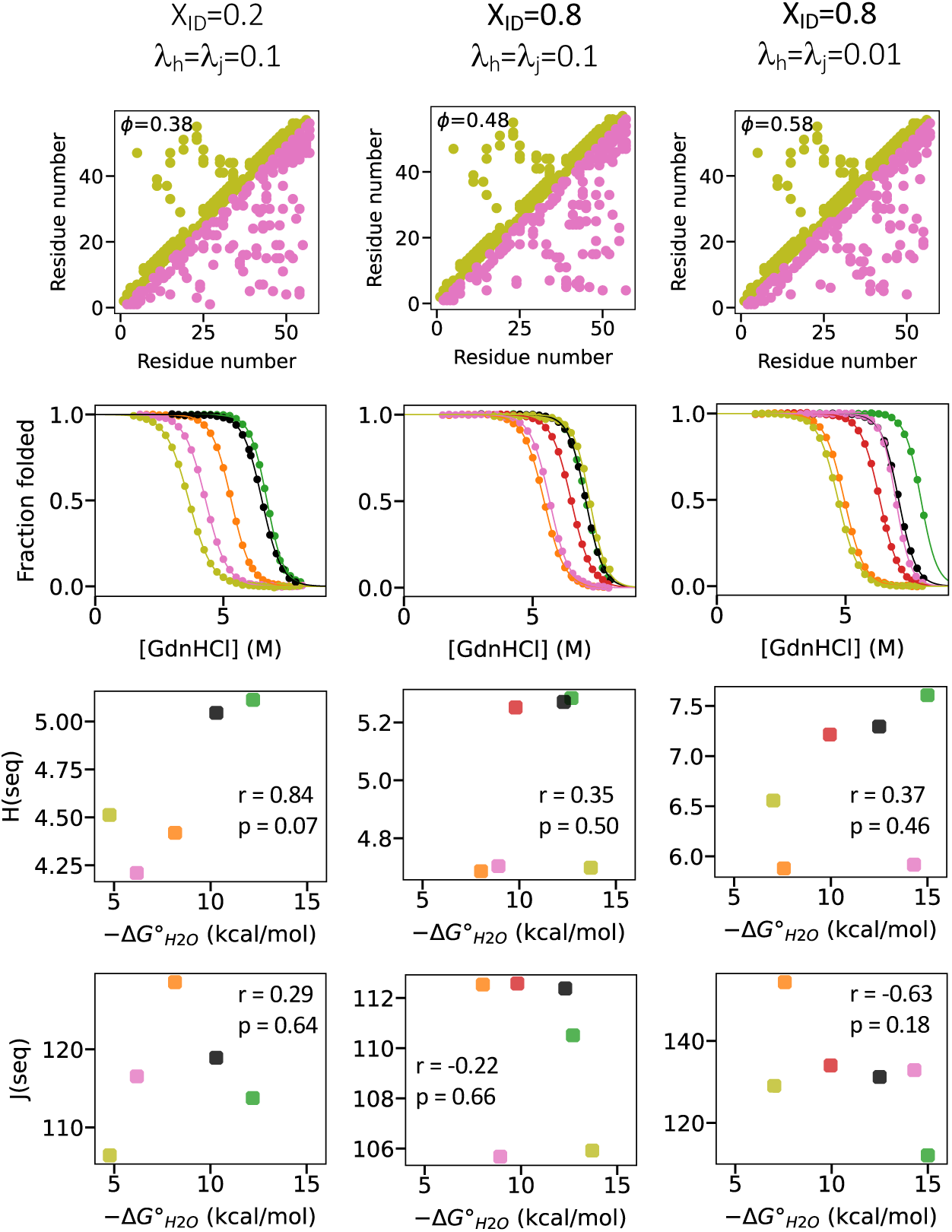
Analysis of Potts model parameterization and designed protein stabilities. The three columns show analyses using single-site and coupling coefficients determined from different fits of the Potts model to the same HD MSA, but using different values of sequence reweighting threshold (X_ID_) and regularization coefficients (λ_j_=λ_h_) when fitting the model. (A) Comparison of top 10% of pair positions closest in structure (yellow, upper triangle) and positions with the top 10% strongest coupling from the Potts model (grey, lower triangle). The Matthews correlation coefficient (ϕ) between the position pairs with the strongest coupling coefficients and closest contacts shown on each plot. (B) GdnHCl-induced unfolding transitions of the single lowest-energy HD sequences for each energy function. All transitions were collected at 25 °C. Solid lines represent the fit of a two-state unfolding model. Proteins too stable to achieve an unfolded baseline at 25 °C were analyzed using a global model of GdnHCl transitions at multiple temperatures (see Methods, Fig S5). All other proteins were analyzed using individual GdnHCl transitions at 25 °C. (C) Correlation between folding free energies and sequence single-site energies determined by Eq 5.2A. (D) Correlation between folding free energies and sequence coupling energies determined by Eq 5.2B. In panels C and D, Pearson correlation coefficients (r) and p-values for correlation coefficients are shown. Colors as in Figure S8. All fitted parameters and sequences shown in Tables S4 and S5.

**Figure S10.**
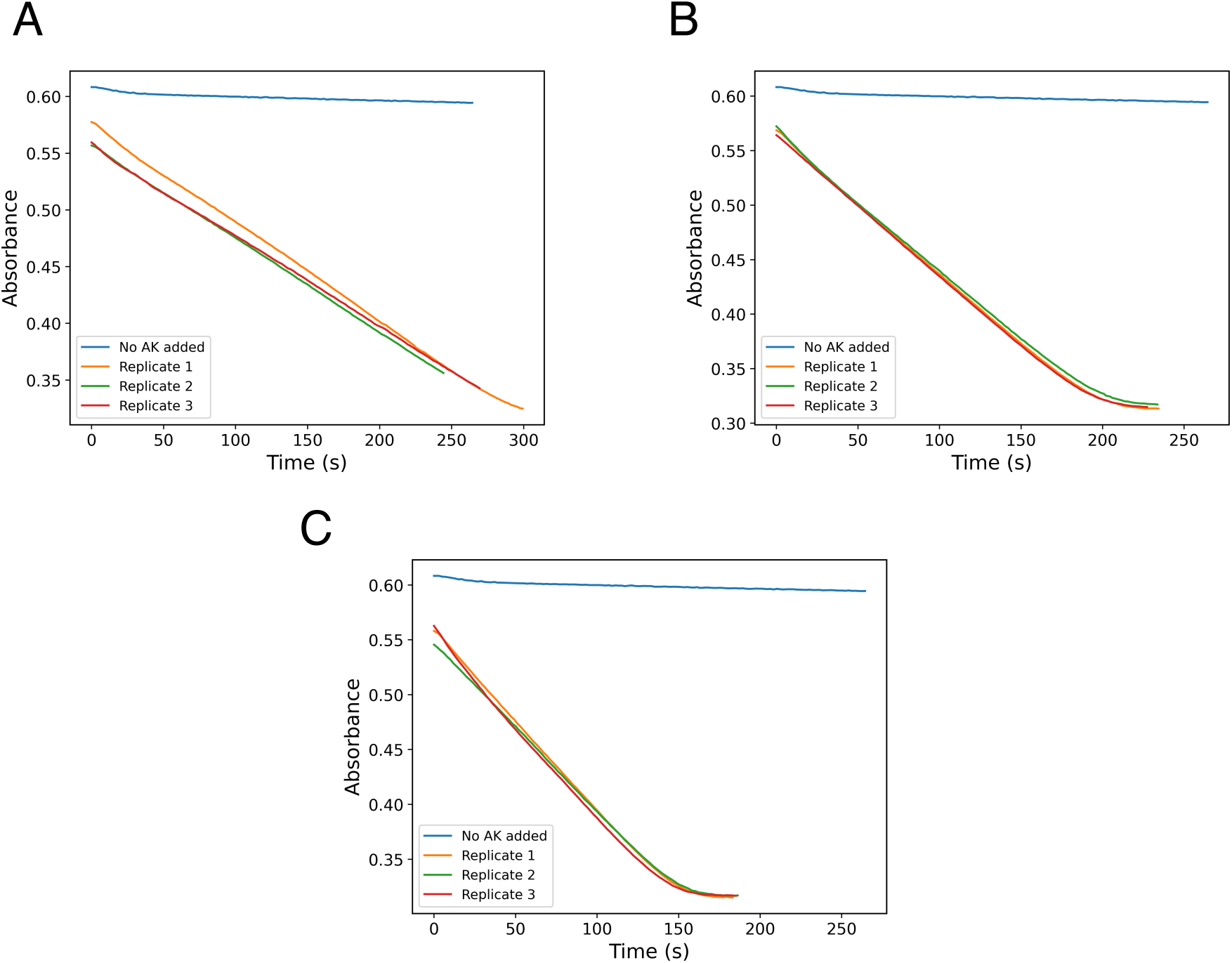
Steady-state enzyme kinetics of designed AKs. Activities of the H-optimized (A), HJ-optimized (B), and consensus (C) AKs were measured using a spectroscopic assay by monitoring the coupled oxidation of NADH by absorbance at 340 nm. Reactions were carried out with AK concentrations of 0.9 nM for the single-site optimized, 0.4 nM for the single-site and coupling optimized, and 122 nM for the consensus. All reactions were at 25 °C.

**Table S1.**
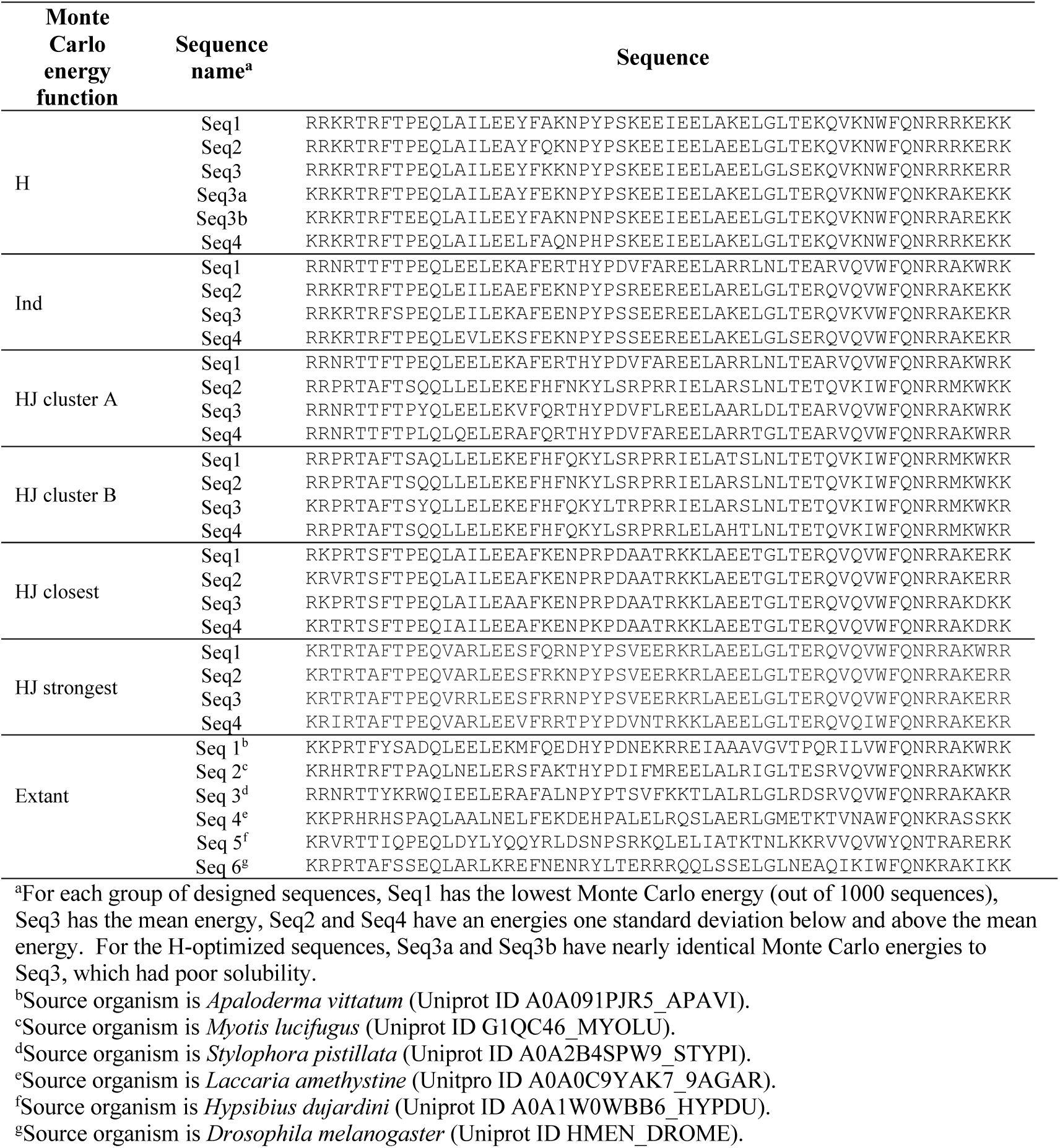
Homeodomain sequences from Potts design with λ_j_=λ_h_=0.01, X_ID_=0.8.

**Table S2.**
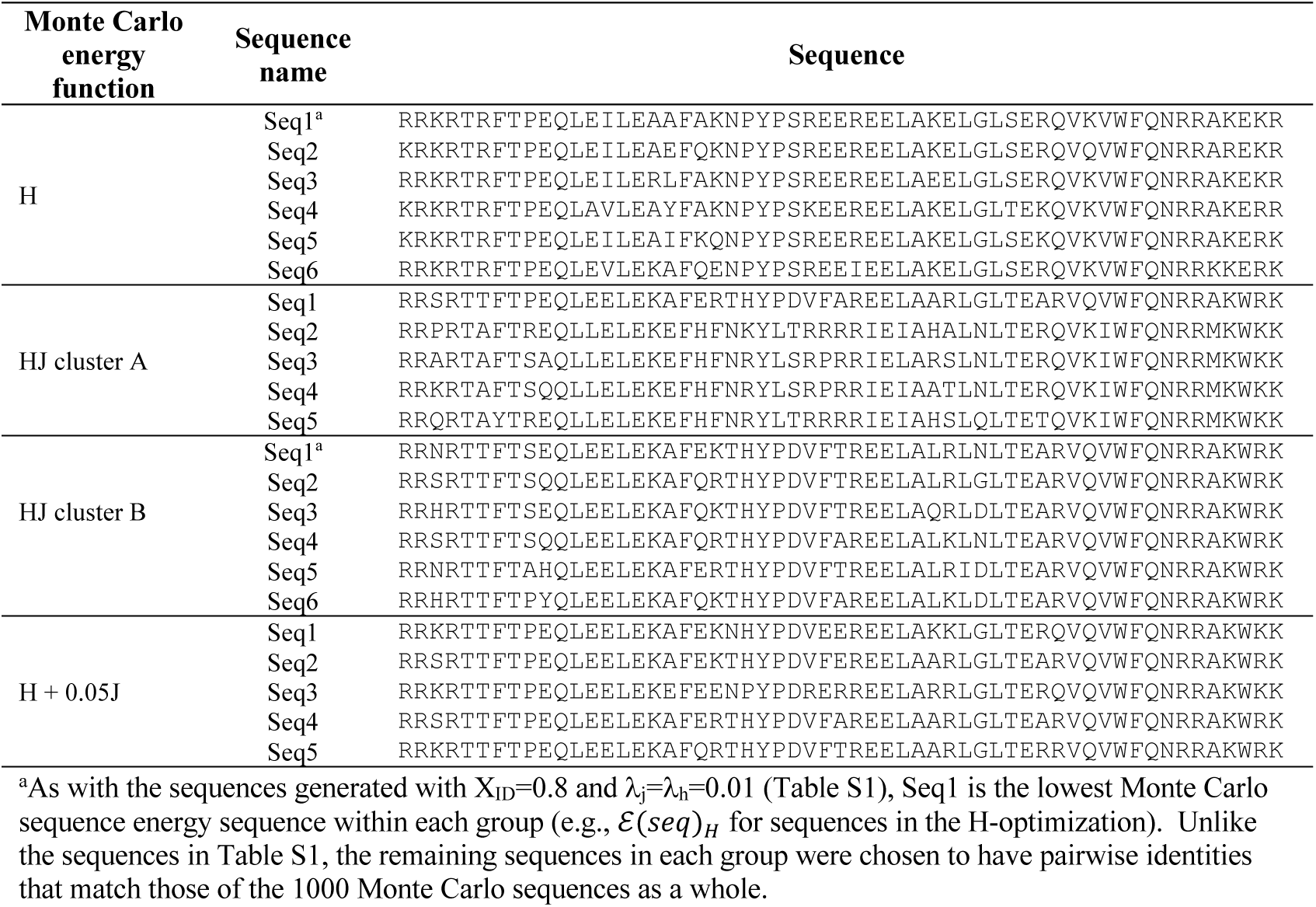
Homeodomain sequences from Potts design with λ_j_=λ_h_=0.1, X_ID_=0.2.

**Table S3.** Unfolding thermodynamics for Potts and extant HDs with λ_j_=λ_h_=0.01, X_ID_=0.8.

| Potts energy function | $\Delta G^{\circ}_{H_2O}$<sup>a</sup><br>(kcal mol<sup>-1</sup>) | m-value<sup>a</sup><br>(kcal mol<sup>-1</sup> M<sup>-1</sup>) | C<sub>m</sub><sup>a</sup><br>(M) |
| --- | --- | --- | --- |
| H Seq1<sup>b</sup> | -15.0 ± 0.3 | 1.89 ± 0.04 | 7.92 ± 0.22 |
| H Seq2<sup>b</sup> | -15.2 ± 1.33 | 1.83 ± 0.18 | 8.30 ± 1.1 |
| H Seq3a | -14.33 ± 0.51 | 1.77 ± 0.07 | 8.10 ± 0.4 |
| H Seq3b | -13.38 ± 0.18 | 1.68 ± 0.02 | 7.96 ± 0.15 |
| H Seq4<sup>b</sup> | -12.0 ± 0.15 | 1.71 ± 0.02 | 7.02 ± 0.12 |
| H mean | -14.0 |  |  |
| HJA_Seq1<sup>c</sup> | -7.48 ± 0.08 | 1.46 ± 0.01 | 5.13 ± 0.04 |
| HJA_Seq2<sup>c</sup> | -9.12 ± 0.06 | 1.56 ± 0.01 | 5.84 ± 0.02 |
| HJA_Seq3<sup>c</sup> | -9.36 ± 0.02 | 1.62 ± 0.01 | 5.76 ± 0.01 |
| HJA_Seq4 | -6.84 ± 0.38 | 1.49 ± 0.07 | 4.57 ± 0.03 |
| HJA mean | -8.2 |  |  |
| HJB_Seq3<sup>c</sup> | -6.45 ± 0.13 | 1.46 ± 0.03 | 4.71 ± 0.01 |
| HJS_Seq1<sup>b</sup> | -12.8 ± 0.23 | 1.75 ± 0.03 | 7.34 ± 0.19 |
| HJS_Seq2<sup>b</sup> | -10.4 ± 0.11 | 1.49 ± 0.02 | 6.97 ± 0.10 |
| HJS_Seq3<sup>b</sup> | -10.9 ± 0.09 | 1.61 ± 0.01 | 6.78 ± 0.08 |
| HJS_Seq4<sup>c</sup> | -10.3 ± 0.18 | 1.61 ± 0.02 | 6.40 ± 0.02 |
| HJS mean | -11.1 |  |  |
| HJC_Seq1<sup>c</sup> | -10.5 ± 0.10 | 1.64 ± 0.02 | 6.42 ± 0.01 |
| HJC_Seq2<sup>c</sup> | -10.5 ± 0.2 | 1.62 ± 0.03 | 6.47 ± 0.02 |
| HJC_Seq3<sup>c</sup> | -10.3 ± 0.05 | 1.63 ± 0.01 | 6.30 ± 0.01 |
| HJC_Seq4<sup>c</sup> | -9.49 ± 0.06 | 1.56 ± 0.01 | 6.09 ± 0.01 |
| HJC mean | -10.2 |  |  |
| Ind_Seq1<sup>b</sup> | -12.5 ± 0.1 | 1.77 ± 0.02 | 7.07 ± 0.12 |
| Ind_Seq2<sup>b</sup> | -11.1 ± 0.17 | 1.56 ± 0.02 | 7.10 ± 0.16 |
| Ind_Seq3<sup>c</sup> | -9.83 ± 0.26 | 1.55 ± 0.04 | 6.33 ± 0.02 |
| Ind_Seq4 | -8.49 ± 0.04 | 1.44 ± 0.01 | 5.90 ± 0.01 |
| Ind mean | -10.5 |  |  |
| Extant Seq1 | -5.32 ± 0.12 | 1.42 ± 0.02 | 3.76 ± 0.01 |
| Extant Seq2 | -5.45 ± 0.28 | 1.52 ± 0.06 | 3.57 ± 0.03 |
| Extant Seq3<sup>d</sup> | ND | ND | ND |
| Extant Seq4 | -2.97 ± 0.03 | 1.44 ± 0.01 | 2.06 ± 0.01 |
| Extant Seq5 | -2.38 ± 0.06 | 1.34 ± 0.02 | 1.77 ± 0.02 |
| Extant Seq6 | -3.44 ± 0.01 | 1.43 ± 0.01 | 2.40 ± 0.01 |
| Extant mean | -3.91 |  |  |

<sup>a</sup>All values reported for unfolding thermodynamics at 25 °C. Unless otherwise noted, values represent the mean from fits of a two-state model to two independent titrations and errors represent standard errors of the mean.
<sup>b</sup>Values determined from a global fit of unfolding transitions at multiple temperatures. Errors represent standard errors determined from the fit.
<sup>c</sup>Values are the mean determined from multiple experiments. Errors are the standard deviation on the mean.
<sup>d</sup>Values for extant Seq3 were not determined since the protein did not remain soluble.

**Table S4.**
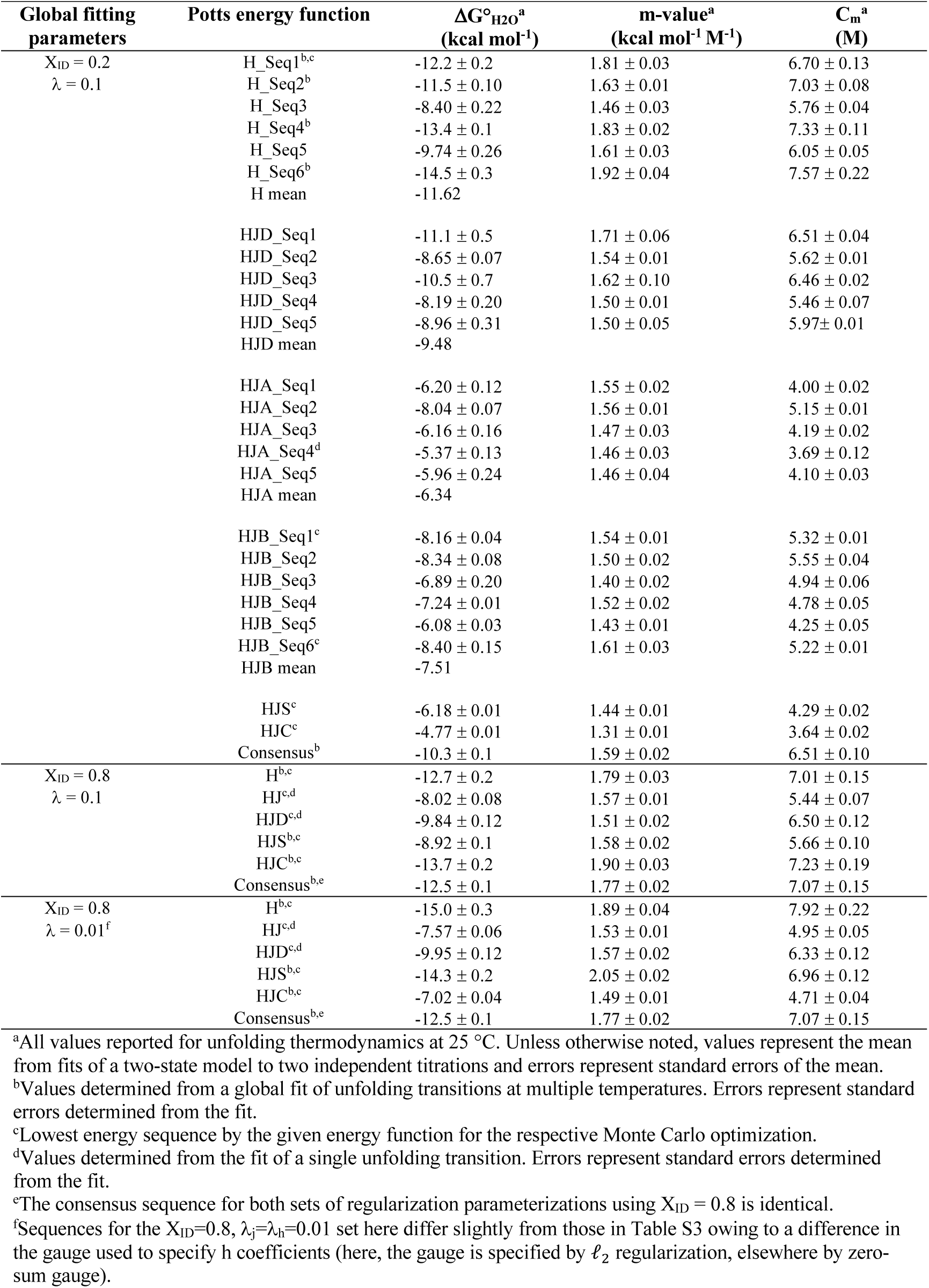
Unfolding thermodynamics for Potts HDs with alternative global fitting parameters.

**Table S5.**
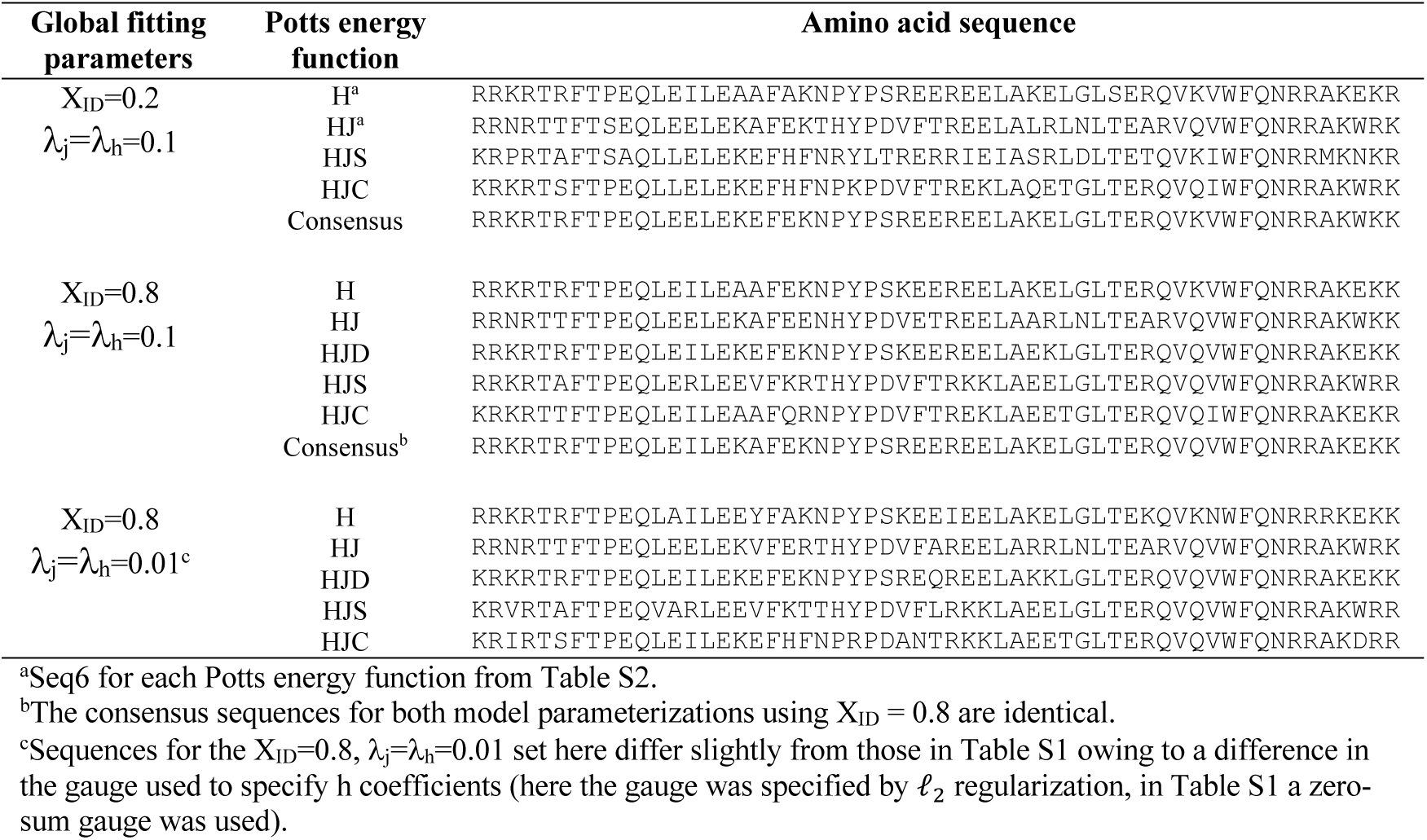
HD lowest energy sequences.

**Table S6.** Comparison of the HD lowest-energy sequences from different generated using different global fitting parameters.

| Potts energy<br>function | $X_{ID}=0.2, \lambda_j=\lambda_h=0.1$ | $X_{ID}=0.2, \lambda_j=\lambda_h=0.1$ | $X_{ID}=0.8, \lambda_j=\lambda_h=0.1$ |
| --- | --- | --- | --- |
| | vs.<br>$X_{ID}=0.8, \lambda_j=\lambda_h=0.1$ | vs.<br>$X_{ID}=0.8, \lambda_j=\lambda_h=0.01$ | vs.<br>$X_{ID}=0.8, \lambda_j=\lambda_h=0.01$ |
| H | 93% | 82% | 86% |
| HJ | 89% | 91% | 88% |
| HJD | 88% | 88% | 93% |
| HJS | 49% | 49% | 89% |
| HJC | 72% | 79% | 77% |
| Consensus | 93% | 93% | 100% |

Values given are percent sequence identities between the lowest energy sequence from each indicated energy function using different values of for the sequence reweighting threshold ( $X_{ID}$ ) and regularization strength ( $\lambda_h, \lambda_j$ ) during Potts optimization.

**Table S7.**
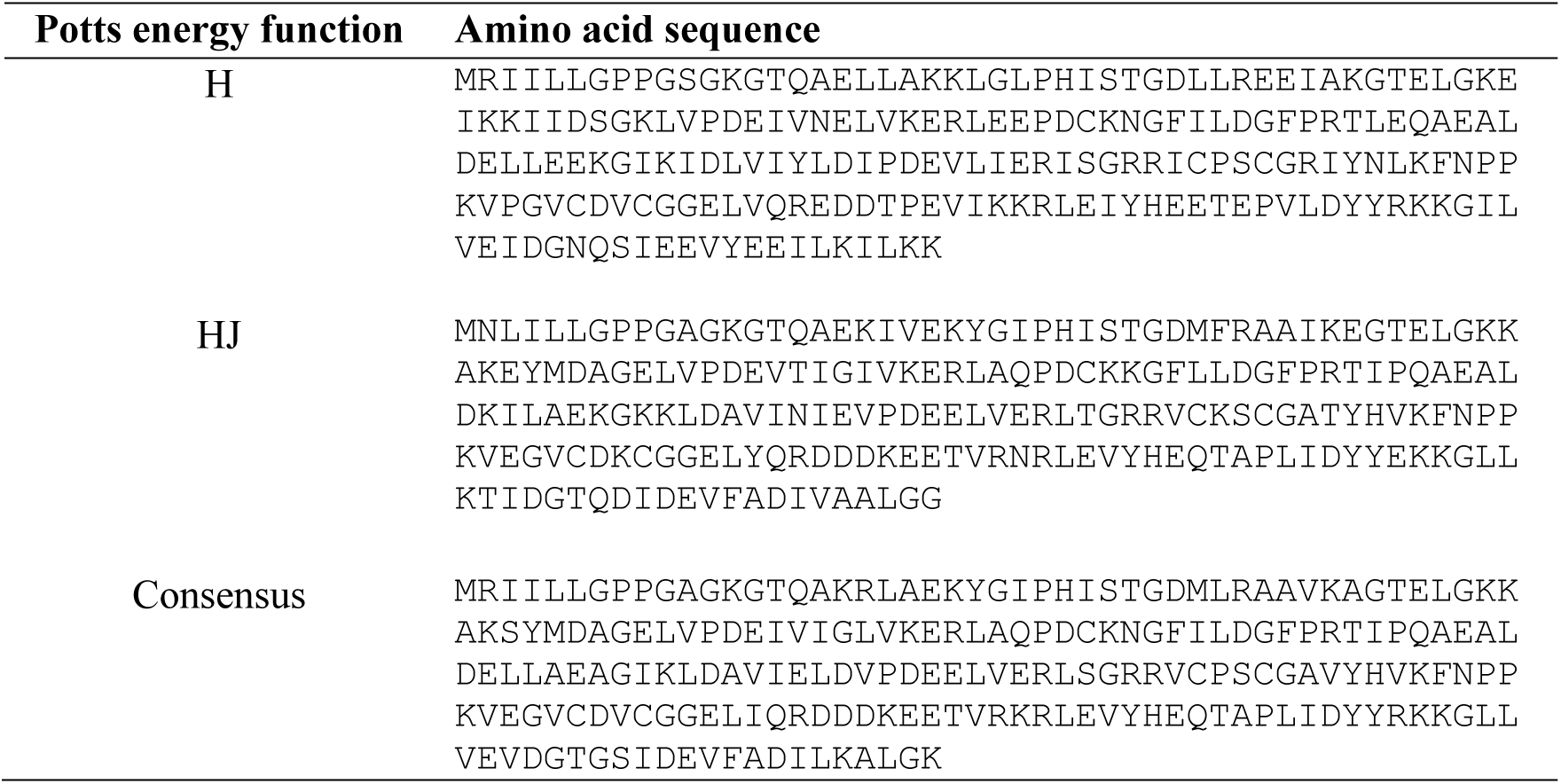
AK designed sequences.

**Table S8.** AK unfolding thermodynamics.

| Potts energy<br>function | $\Delta G^{\circ}_{H_2O}$<sup>a</sup><br>(kcal mol<sup>-1</sup>) | m-value<sup>a</sup><br>(kcal mol<sup>-1</sup> M<sup>-1</sup>) | $C_m$<sup>a</sup><br>(M) |
| --- | --- | --- | --- |
| H | $-17.9 \pm 1.0$ | $3.25 \pm 0.18$ | $5.50 \pm 0.40$ |
| H + J | $-11.2 \pm 0.6$ | $3.62 \pm 0.17$ | $3.11 \pm 0.21$ |
| Consensus | $-18.1 \pm 0.5$ | $5.67 \pm 0.12$ | $3.87 \pm 0.15$ |

<sup>a</sup>Values determined from the fit of a single unfolding transition to a two-state model. Errors represent standard errors determined from the fit. All unfolding transitions were determined at 20 °C.

**Table S9.** HD DNA binding thermodynamics.

| Protein | $K_d$<br>(nM) | $\Delta H$<br>(kcal mol<sup>-1</sup>) | $-T\Delta S$<br>(kcal mol<sup>-1</sup>) |
| --- | --- | --- | --- |
| H Seq1<sup>a</sup> | $8.9 \pm 1.3$ | $-11.1 \pm 0.09$ | $0.35 \pm 0.13$ |
| Ind Seq1<sup>a</sup> | $4.2 \pm 0.9$ | $-14.0 \pm 0.1$ | $2.8 \pm 0.14$ |
| HJA Seq1<sup>a</sup> | $340 \pm 30$ | $-8.8 \pm 0.15$ | $0.12 \pm 0.21$ |
| H Seq1<sup>b</sup> | $7.7 \pm 1.3$ | $-15.4 \pm 0.1$ | $4.5 \pm 0.1$ |
| HJB Seq1<sup>b</sup> | $110 \pm 12$ | $-10.1 \pm 0.1$ | $0.77 \pm 0.15$ |
| Consensus<sup>d</sup> | $8.1 \pm 1.6$ | $-16.0 \pm 0.2$ | $5.0 \pm 0.3$ |
| *D. melanogaster*<br>Engrailed<sup>d</sup> | $737 \pm 38$ | $-9.2 \pm 0.1$ | $0.8 \pm 0.1$ |

<sup>a</sup>Proteins are the lowest-energy sequences from optimizations using coefficients from the $X_{ID} = 0.8$ , $\lambda_j = \lambda_h = 0.01$ model parameterization.
<sup>b</sup>Proteins are the lowest-energy sequences from optimizations using coefficients from the $X_{ID} = 0.2$ , $\lambda_j = \lambda_h = 0.1$ model parameterization.
<sup>c</sup>Values are determined from the fit of a single titration to a one-site binding model. Errors represent standard errors determined from the fit.
<sup>d</sup>Values for Consensus are from Tripp et al.<sup>18</sup> This consensus differs from the consensus HDs in this study by 8 residues ( $X_{ID} = 0.2$ ) and 5 residues ( $X_{ID} = 0.8$ ) owing to differences in the MSAs and reweighting parameters.

**Table S10.**
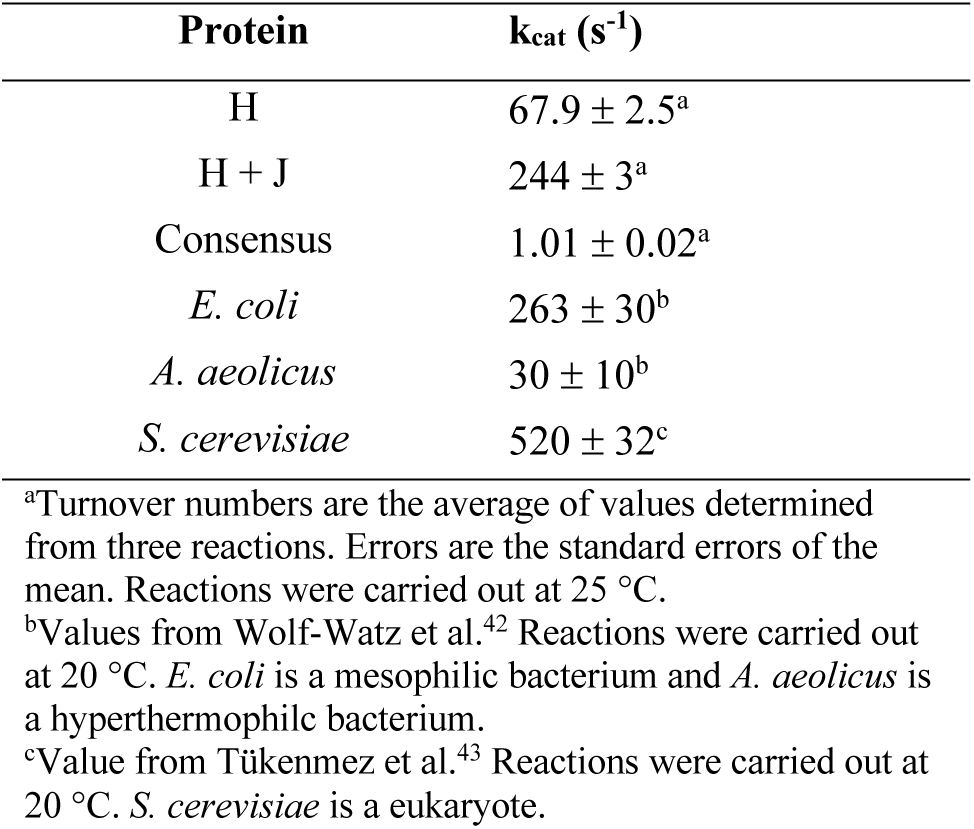
AK steady-state turnover numbers.

## Notes

### Competing Interest Statement

The authors have declared no competing interest.

